# Systematic Analysis of Biological Processes Reveals Gene Co-expression Modules Driving Pathway Dysregulation in Alzheimer’s Disease

**DOI:** 10.1101/2024.03.15.585267

**Authors:** Temitope Adeoye, Syed I Shah, Ghanim Ullah

## Abstract

Alzheimer’s disease (AD) manifests as a complex systems pathology with intricate interplay among various genes and biological processes. Traditional differential gene expression (DEG) analysis, while commonly employed to characterize AD-driven perturbations, does not sufficiently capture the full spectrum of underlying biological processes. Utilizing single-nucleus RNA-sequencing data from postmortem brain samples across key regions—middle temporal gyrus, superior frontal gyrus, and entorhinal cortex—we provide a comprehensive systematic analysis of disrupted processes in AD. We go beyond the DEG-centric analysis by integrating pathway activity analysis with weighted gene co-expression patterns to comprehensively map gene interconnectivity, identifying region- and cell-type specific drivers of biological processes associated with AD. Our analysis reveals profound modular heterogeneity in neurons and glia as well as extensive AD-related functional disruptions. Co-expression networks highlighted the extended involvement of astrocytes and microglia in biological processes beyond neuroinflammation, such as calcium homeostasis, glutamate regulation, lipid metabolism, vesicle-mediated transport, and TOR signaling. We find limited representation of DEGs within dysregulated pathways across neurons and glial cells, indicating that differential gene expression alone may not adequately represent the disease complexity. Further dissection of inferred gene modules revealed distinct dynamics of hub DEGs in neurons versus glia, highlighting the differential impact of DEGs on neurons compared to glial cells in driving modular dysregulations underlying perturbed biological processes. Interestingly, we note an overall downregulation of both astrocyte and microglia modules in AD across all brain regions, suggesting a prevailing trend of functional repression in glial cells across these regions. Notable genes, including those of the CALM and HSP90 family genes emerged as hub genes across neuronal modules in all brain regions, indicating conserved roles as drivers of synaptic dysfunction in AD. Our findings demonstrate the importance of an integrated, systems oriented approach combining pathway and network analysis for a comprehensive understanding of the cell-type-specific roles of genes in AD-related biological processes.

## Background

Alzheimer’s disease (AD) is an increasingly prevalent neurodegenerative disorder with global cases surpassing 50 million, presenting an urgent need for understanding its complex pathology (1,2). The etiology of AD is characterized by hallmark molecular and cellular alterations, most notably the accumulation of senile amyloid-beta (Aβ) plaques and the presence of hyperphosphorylated Tau neurofibrillary tangles (NFTs) (2–6). Such pathological alterations often trigger neurotoxic cascades, resulting in synaptic dysfunction, pervasive neuronal loss, and subsequent functional disruption of neuronal networks (7–11). However, AD perturbations manifest heterogeneously across brain regions and cell types, contributing to the complexity of its pathology (12,13). Indeed, several lines of evidence indicate that AD inflicts selective disruptions to biological processes among cellular subpopulations in different brain regions, revealing a region- and cell-type-dependent susceptibility (14–17). This cellular and regional diversity in affected mechanisms poses significant challenges in the discovery and screening of candidate biomarkers and potential therapeutic strategies.

Recent advancements in single-cell/single-nucleus RNA-sequencing (sc/snRNA-seq) present an opportunity to dissect the molecular basis of AD with unprecedented resolution (18,19). Leveraging these techniques, numerous studies have identified differential gene expression (DEG) patterns associated with AD, revealing insights into cellular states and their variations during disease progression (20–23). For instance, gene expression analysis of cells in the prefrontal cortex revealed that neurons primarily contain downregulated genes in AD, while glial cells, albeit to a lesser extent, exhibit opposite directionality (21). Indeed, top DEGs were cell type-specific, highlighting the distinct cell-type-specific transcriptional responses to AD-associated perturbations. Consistent with this, Grubman et al.(24) identified upregulated transcription factors in the entorhinal cortex that mediate cell-type-specific state transitions from control to AD. Similarly, comprehensive transcriptomic evaluations in human and mice models revealed a unique set of DEGs associated with a disease-associated microglia (DAM) state (22,23,25). Notably, these studies revealed that the DAM state is marked by downregulation of several homeostatic genes, recapitulating the notion that cell-type subpopulations can express distinct transcriptional alterations. Recently, Habib et al. (26) reported an AD-associated astrocyte subpopulation in the prefrontal cortex and hippocampus, characterized by elevated GFAP levels and increased expression of genes implicated in amyloid aggregation and inflammation (22,26). Despite the detailed transcriptional landscape of AD outlined by these findings, such investigations predominantly focus on isolated differential gene expressions, lacking an integrated systems-level understanding of the relationships between these genes and their functions within broader biological processes.

AD is recognized as a systems disease, where the pathology extends beyond molecular alterations to encompass complex interactions in gene networks (27,28). The pathological progression and perturbation of biological processes in AD are not merely driven in isolation by DEGs, but rather by the complex interplay of a robust sets of genes within biological processes or signaling cascades (29). Thus, the collective molecular interactions observed across various cellular processes fundamentally shape the pathogenesis of AD (30). Gene co-expression network analyses have emerged as critical tools to capture these interactions, uncovering highly interconnected network of genes in AD and higher order network structures associated with the pathology. Notably, Morabito et al. (31) utilized this systems-level perspective to identify consensus networks of microglia genes representing classical markers of homeostatic microglia or known DAM genes, indicating that microglia assume activated states due to the functional interplay of associated genes. Likewise, Miyoshi et al. (32) demonstrated unique dysregulation patterns in functional biological units in early sporadic AD, suggesting that dynamic modular changes in gene expression may play a crucial role in AD progression. These findings collectively offer a thorough characterization of the systems-level features of the AD brain. However, since functional perturbation of biological processes arises from the underlying network architecture of the comprising gene programs, it is still unclear whether and to what extent DEGs play a central role in the perturbation of these processes or whether they are merely partakers of their associated biological units (33,34).

In this study, we leverage snRNA-seq data from key regions of postmortem AD brains to conduct a systems-level analysis of pathway perturbations. Our approach integrates pathway activity analysis with weighted gene co-expression patterns, providing insights into functional coherence and interplay among genes involved in perturbed biological processes in AD. To identify the complex systems-level changes in both neuronal and glial cell populations, we first comprehensively characterize region- and cell-type-specific pathway dysregulation patterns associated with AD. This nuanced approach reveals an expanded role for astrocytes and microglia in a variety of biological processes than previously appreciated in neuron-centric models of AD. We also highlight the dysregulation of calcium (Ca^2+^) signaling across different cell types and regions, representing an axis of disruption that has been consistently implicated in AD pathology. Next, we qualitatively demonstrate that DEGs are not robustly distributed in the curated set of genes comprising the biological processes (gene programs) implicated in AD. Finally, we employ a weighted gene co-expression strategy to uncover gene modules and highly connected hub genes underlying the perturbed gene programs. This approach revealed distinct dynamics of hub DEGs (hub-DEGs) in neuronal versus glial modules, which suggests that DEGs exert a more pronounced influence on neurons than on glial cells in driving pathway perturbations in AD. By offering a comprehensive, systems-driven perspective of AD pathology, our findings refine the current understanding of the disease and opens new avenues for targeted therapeutic and diagnostic strategies.

## Methods

### Study Design and Data Acquisition

Here, we leveraged pre-processed snRNA-seq data obtained from two independent studies (20,35) comprising three different brain regions: Middle Temporal Gyrus (MTG), Superior Frontal Gyrus (SFG), and Entorhinal Cortex (ETC) (*Fig. 1*). We reasoned that using samples matched for pathological status would minimize the technical variation due to data composition and allow for meaningful comparison across the brain regions. To accomplish this, we selected a cohort of 10 male individuals form the Gabitto et al. (MTG) (35) study based on their level of AD neuropathologic change (ADNC). Donors from the Gabitto et al. study (35) were specifically chosen to align with corresponding cases from the Leng et al. (SFG & ETC) study (20). ADNC stage is evaluated using the robust “ABC” scoring system, considering the Thal phases (A) to gauge the overall Aβ burden; Braak stage (B) for neurofibrillary tangles (NFT) load, and neuritic plaque score (C) (36). The combination of A, B, and C scores are used to categorize individuals into distinct pathological stages, denoted as “Not AD”, “Low”, “Intermediate” and “High” ADNC stages. It is worth mentioning that accumulation of Aβ plaques and extent of NFT inclusions have consistently proven to be the most reliable correlates of neuropathological staging and AD diagnosis (37–39). Consequently, “Intermediate” or “High” AD neuropathologic stages are typically associated with dementia. To ensure a balanced representation across ADNC stages, each study cohort comprised four individuals with a “Not AD” descriptor, representing cognitively healthy controls, while the remaining six were equally distributed among Low/Intermediate and High ADNC stages, allowing us to capture the cortical-free, early, and late stages of AD pathology in our analysis (*Fig. 1*). The pre-processed data (obtained after quality-control filtering) from (35) contained 154,368 snRNA-seq profiles from the MTG, while a total of 106,136 nuclei (SFG = 63,608 & ETC = 42,528) were obtained from (20) (*Fig. 1*). Predefined cell-type annotations were used to restrict analysis to six different cell types: excitatory neurons, inhibitory neurons, astrocytes, microglia, oligodendrocytes, and oligodendrocyte precursor cells (OPCs).

**Fig. 1.**
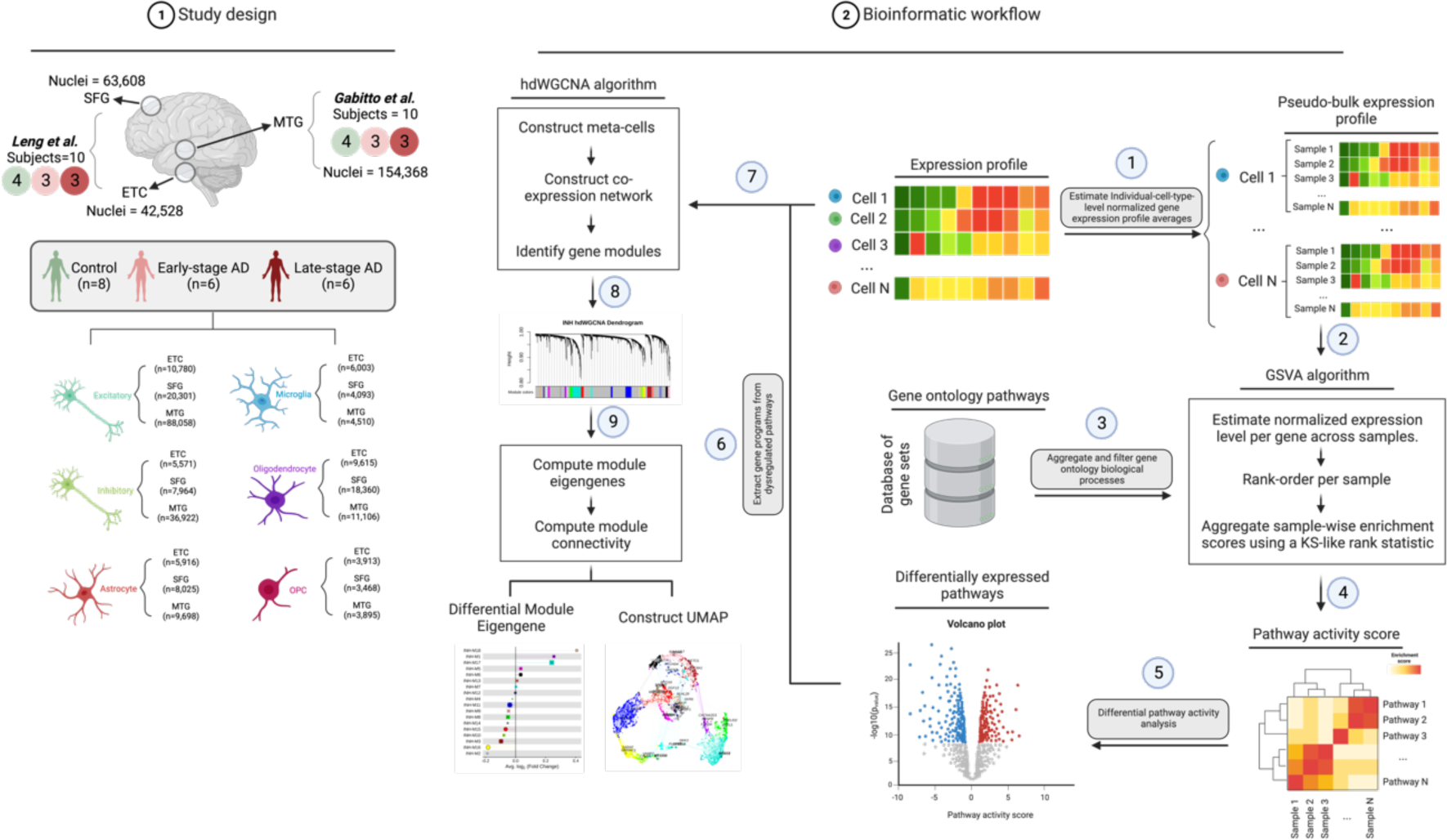
Schematic overview of study workflow and analytical methods. Sample and nuclei distribution across pathological and study groups are shown in the left panel, while the right panel illustrates the bioinformatics pipeline employed for the identification of perturbed gene ontology (GO) biological processes and their associated gene co-expression networks (**Methods**), created using <u>BioRender.com</u>. The workflow begins with computing pseudo-bulked averages of normalized gene expression profiles for single-cell expression profiles of each cell type (Steps 1 & 2). Subsequently, pathway activity scores were calculated (Steps 3 & 4), using gene set variation analysis (GSVA) as previously described (41). Differential pathway activity was then estimated for each pathway-cell type combination, employing a multivariate linear model (Step 5). Construction of co-expression networks, performed with hdWGCNA (31) was specifically limited to gene programs comprising perturbed pathways (Steps 6—8).

The overall cohort of 20 male individuals were originally enrolled in the Adult Changes in Thought (ACT) Study, the University of Washington Alzheimer’s Disease Research Center (ADRC) (35), the Neurodegenerative Disease Brain Bank (NDBB) at UCSF, or the BBAS from the University of Sao Paulo (20,40). These individuals were part of a larger cohort previously reported in (20,35). Notably, brain specimens from the Leng et al. study (20) were obtained from NDBB and BBAS, representing 10 of the male participants from the postmortem cohort used in this study. Brain slices were obtained from ETC and SFG (Brodmann area 8). All individuals underwent rigorous neuropathological assessments following established protocols, ensuring that selected brain samples exhibited pronounced AD-type pathology while excluding non-AD pathologies, such as Lewy body disease, TDP-43 proteinopathies, primary tauopathies, and cerebrovascular changes.

The study cohort comprised 10 male participants subjected to snRNA-seq, presenting a diverse spectrum of Braak stages (ranging from 0 to 6), ADNC categories (comprising Not AD [n=4], Low [n=3], and High [n=3]), and consistently harboring APOE ε3/ε3 genotypes. The isolation of nuclei was extensively documented in (20). Briefly, postmortem frozen brain tissue was dounce-homogenized with the addition of IGEPAL-630, followed by gradient centrifugation for nuclei filtration and purification. Subsequently, sequencing libraries were constructed utilizing droplet-based snRNA-seq with 10X Genomics’ Chromium Single Cell 3’ Reagent Kits v2, targeting a total of 10,000 nuclei per sample. The resulting sequencing data underwent demultiplexing through Cell Ranger, utilizing a customized pre-mRNA GRCh38 reference genome designed to accommodate introns. Alignment and gene expression quantification were performed using cellranger count under default settings.

Brain specimens from the study by Gabitto et al. (35) were obtained from the ACT Study and the UW ADRC. The study cohort was carefully selected, encompassing a wide spectrum of AD severity while excluding individuals diagnosed with Frontotemporal Dementia, Frontotemporal Lobar Degeneration, Down Syndrome, Amyotrophic Lateral Sclerosis, or other degenerative disorders (except Lewy Body Disease). The cohort consisted of 84 participants aged 65 and above, representing various stages of Alzheimer’s disease severity. Rapid autopsies were conducted to ensure a postmortem interval of less than 12 hours. Tissue processing involved uniform coronal slicing of one hemisphere, fixed or frozen slabs, and subsequent processing of Superior and Middle Temporal Gyrus tissue samples. As in (20), the study rigorously adhered to neuropathological assessments, tissue processing, and immunohistochemical analyses, providing clinical, cognitive, and demographic data. Specifically, the study comprised a cohort of 84 ACT/ADRC donors spanning a broad range of ADNC levels and comorbid pathologies, including Lewy Body Disease, vascular brain injury, and hippocampal sclerosis. Notably, the cohort tended to skew towards more advanced stages of the disease, with 58% of participants exhibiting a Braak stage of 5 or higher (Braak Stage: 0 [n=2], 2 [n=4], 3 [n=6], 4 [n=23], 5 [n=34], 6 [n=15]) and 61% having a Thal Phase of 4 or higher (Thal Phase: 0 [n=9], 1 [n=5], 2 [n=7], 3 [n=12], 4 [n=30], 5 [n=21]). Demographically, the cohort displayed a slight female bias (51 females and 33 males), particularly among donors with high ADNC (Not AD [N=9], Low [n=12], Intermediate [n=21], High [n=42]). Furthermore, the cohort was characterized by advanced age, with an average age at death of 88 years, and half of the donors received a clinical diagnosis of dementia. Genetic analysis revealed the presence of the APOE ε4 genotype, a primary risk factor for AD, in 23 donors, while the remaining donors possessed ε3 and ε2 alleles in various combinations. Sequencing libraries were constructed following standard guidelines for 10x Genomics kits. RNA isolation from nuclei was performed with subsequent evaluation of RNA integrity. The isolated nuclei and high-quality RNA samples were then employed for snRNA-seq, snATAC-seq, and MERFISH. The selection of individuals from this study for our work was based on their alignment with corresponding cases from the participants in (20), considering their level of ADNC. For the present analysis, participants were stratified based on the ADNC spectrum, encompassing individuals from “Not AD” to various stages of AD pathology as outlined above.

### Data Processing

The processed droplet-based snRNA-seq profiles, amounting to a total of 260,504 profiles, were obtained from (42,43). Quality control and filtering steps were previously detailed in each study. In Leng et al. (20), raw gene-barcode matrices were converted into SingleCellExperiment (SCE) objects in R using DropletUtils. Nuclei from empty droplets or with fewer than 200 UMIs were discarded, followed by data merging and normalization based on the strategy in (44). High-variance genes were identified for dimensionality reduction using the Seurat package, but as individual origin influenced results, the scAlign tool was adopted for cross-sample alignment, prioritizing biological over technical factors. Clusters were mapped to major brain cell types using specific marker genes, with ambiguous clusters removed, and fine-grained subclustering performed by isolating cells from primary cell types. In the study Gabitto et al. (35), nuclei gene expression data were mapped to a reference transcriptome using gene expression and chromatin accessibility profiles, discarding nuclei with fewer than 500 detected genes from upstream of cell type mapping. The filtered nuclei were then classified into classes, subclasses, and supertypes using scANVI, with predictions evaluated against known marker gene expressions. Regions with variable expression were examined for potential contamination, and data were further refined using high-resolution Leiden clustering. Clusters with undesirable metrics were subsequently flagged and removed to further improve quality.

The samples from the ETC and SFG of autopsied brains generated by Leng et al. (20) is accessible for download from Synapse.org (42) under the Synapse ID syn21788402. Gabitto et al. (35) data, generated from the MTG, was obtained from the Seattle Alzheimer’s Disease (SEA-AD) Brain Cell portal (43,45). We categorized participants into three distinct groups: 8 individuals with “Not AD” designation served as cognitively healthy controls, while 12 individuals manifested mild to severe AD-pathology. Out of these 12 AD-pathology group, 6 participants with “Low” or “Intermediate” ADNC scores are designated as ’early-pathology’ group, whereas the remaining 6 with a “High” ADNC scores are designated as ’late-pathology’ group. As reported in the source studies, informed consent was obtained for all participants, and ethical approvals for the use of human tissues were obtained from the respective institutional review boards. All post-mortem neuropathological assessments, clinical evaluations, and pathological grouping are detailed in Supplementary Table 1.

### Differential gene expression

Cell type-specific differential gene expression analysis was evaluated using a customized version of the Libra R package (46) accessible via GitHub. The source package implements 22 unique differential expression methods that can all be accessed from a singular function call. Given the susceptibility of cell-based differential expression methods to the drop-out events and overdispersion intrinsic to single-cell data, we mitigated against these limitations by using a method designed specifically for bulk sequencing data. Specifically, we adopted the DESeq2 (47) routine with the Wald test for differential expression analysis between the control group and the AD group. To ensure that the analysis accounted for true biological replication—that is variability at the level of individual objects—unique molecular identifier (UMI) counts from cells belonging to the same individual and specific cell type were aggregated to create ’pseudo-bulk’ samples. Genes with negligible expression in a given cell type, indicated by a nonzero detection rate below 10% in the aggregated pseudo-bulk, were precluded from further analyses to mitigate false-positive discoveries. Preliminary assessment of the principal components of these individual-level aggregated gene expression profiles corroborated the decision to exclude additional covariates, such as age at death and post-mortem interval. Therefore, the pathological status served as the sole covariate in our differential expression model. Genes were identified as significantly differentially expressed if they exhibited an absolute log fold change exceeding 0.25 and a false discovery rate (FDR) below 0.01. The table of p-values and log fold changes for all genes across all brain regions and cell types is provided in Supplementary Tables 2—4.

### Pathway analyses

The compendium of Gene Ontology biological processes (2018 edition) was retrieved from the Mayaan laboratory repository (48). Certain pathways were renamed to optimize clarity and standard nomenclature, with specific modifications enumerated in Supplementary Table 5.

Pathway activity scores were computed in accordance with protocols outlined in (49). This method effectively retrieved cell type-specific signatures, not accounted for by randomly sampled gene set enrichment analysis (50). In brief, we first computed cell-type-level normalized gene expression profiles for each individual using the ACTIONet normalization procedure (51). Subsequently, pathway activity scores were computed as previously implemented in the R package GSVA (version 1.46.0) (41). GSVA executed with the following parameters: mx.diff=TRUE, kcdf=c(“Gaussian”), min.sz=5, max.sz=500. To minimize the discovery of false positive, gene sets were filtered to exclude genes with insufficient expression in the designated cell type, defined by a nonzero detection rate less than 10%. For each pathway and cell type, activity scores were modeled using a multivariate linear regression, taking the form: activity score ∼*β0 × pathology.group*. No additional covariates were incorporated, as PCA revealed no significant association with pathological status, thus not accounting for observed variances in overall gene profiles. The “pathology.group” variable stratifies samples into ’no-pathology,’ ’early-pathology,’ or ’late-pathology’ categories. Linear models were fitted using the lmfit() function, and corresponding t-statistics were generated through the eBayes() function, both from the Limma R package (version 3.50.3). Differential expression between the ‘no-pathology’ and ‘AD-pathology’ groups was estimated by setting the contrast argument as *makeContrast = (early + late)/2 – no*. Pathways were identified as significantly differentially expressed based on a nominal p-value cut-off of 0.05 (as depicted in Figure 1). The procedure resulted in the identification of prioritized candidate pathways across major cell types. Estimates of β0 coefficients, along with additional statistics as outlined in Figure 1, are comprehensively documented in Supplementary Tables 6—8, including both nominal p-values and FDR-corrected p-values.

### Gene Co-expression Network Analysis

#### Network construction and module identification

To generate robust gene-gene correlations, we employed hdWGCNA (version 0.2.18) (31), specifically tailored for single-cell and scRNA-seq data. We first generated a Seurat object (version 4.3.0.1) (52) using the ‘SetupForWGCNÀ function, setting the “gene_select” parameter to custom. We confined our analysis to functionally relevant gene programs, extracted from the gene sets comprising the pathways that were dysregulated (with nominal p-values less than 0.05) in each cell type. Metacells, which are essentially aggregates of transcriptionally similar cells originating from the same biological replicate, were constructed using the k-Nearest Neighbors (KNN) algorithm, with default parameters (k=25, max_shared=10). This step mitigated data sparsity inherent to scRNA-seq data and generated a metacell gene expression matrix conducive for robust network construction. Subsequently, the optimal soft power threshold was determined using TestSoftPowers function in a ‘signed’ network, conducting a parameter sweep over a range of 1 to 30. This specifies the degree to which gene-gene correlations are scaled in order to reduce the amount of noise present in the correlation matrix and prioritize strong correlations. The selected soft power thresholds, demonstrating a fit to the scale-free topology model, are reported in Supplementary Figures 3—5. Network construction and module detection were performed using the ConstructNetwork function, which employs the scaled correlations to compute a topological overlap matrix (TOM), reflecting the network of shared neighbors between genes. Module dendrograms were visualized using the PlotDendrogram function (Supplementary Figures 6—8).

### Module signatures and hub gene identification

To summarize the gene signatures within each module, module eigengenes (ME) were calculated using the ModuleEigengenes function with default settings. This effectively represents the first principal components of the subset of the gene expression matrix comprising each module, allowing us to obtain the module feature genes. The intra-modular connectivity (kME), a metric representing the correlation of each gene with its ME, was determined using the SignedKME algorithm, essentially determining the highly connected genes in each module.

### Network visualization

For a comprehensive low-dimensional visualization, we applied the RunModuleUMAP function on the TOM, confining it to the top 5 hub genes per module based on kME values. This resulted in a UMAP representation where the organization was primarily determined by the hub genes, and only the top 10 hub genes in each module were annotated in the UMAP space.

### Differential module analysis and functional enrichment

To discern modular differences between control and diseased group in each cell type, a differential module eigengene analysis was performed using the FindAllMarkers function in Seurat, applying the Wilcoxon test. Results are depicted in lollipop diagrams, with non-significant modules marked “X” (nominal P > 0.05). Supplementary Tables 9—11 contains additional statistics for each cell type across tested brain regions. Furthermore, the overlap of co-expression modules with DEGs or AD-associated genes from the Open Targets Platform (53), KEGG Alzheimer’s disease pathways (54), and Harmonizome (55) was calculated using the R package GeneOverlap (version 1.34.0) via Fisher’s exact test. Finally, functional enrichment analysis was conducted on hdWGCNA modules using the R package enrichR, focusing on Gene Ontology processes exhibiting differential expression in specific cell types.

## Results

### Cell-type- and region-specific perturbations in molecular processes define the heterogeneous responses to AD

To comprehensively characterize perturbed molecular processes in AD, we performed differential pathway activity analysis, leveraging Gene Ontology biological processes (*Fig. 1*, **Methods**). To enhance the sensitivity in detecting subtle changes in pathway activity, we first aggregated gene expression values into pathway activity scores (*Fig. 1*, **Methods**) (49). These scores effectively summarize the collective gene expression levels within each pathway and improved statistical power for subsequent analyses. We then examine whether there are qualitative changes in the aggregated scores due to AD using a multivariate linear model with pathological status as the only covariate. Preliminary analysis of the principal components of the aggregated expression data revealed that other covariates, such as age at death and post-mortem interval, are not correlated with biological or technical variation (*Supplementary Fig. 1*), and as a result were excluded from the design matrix, ensuring that the data modeling focused solely on the biologically relevant factors (56). Pathology groups were defined based on the ADNC levels (Supplementary Table 1, **Methods**), with individuals categorized as early-pathology (low or intermediate ADNC) or late-pathology (high ADNC). These two groups correspond to the pathological progression of AD. early-pathology individuals have discernible amyloid load coupled with mild neurofibrillary tangles and cognitive deficit. Conversely, the late-pathology individuals show higher amyloid burden, elevated NFT deposits, pervasive pathology, and pronounced cognitive impairment (37–39). Both pathology groups were combined in the contrast analysis to assess differential expression between AD-pathology and control groups (**Methods**).

Our analysis revealed that AD inflicts a wide range of perturbations in molecular process (P < 0.05) across the three brain regions (*Fig. 2—4*), ranging from cell-type-specific alterations, exclusive to a single cell type (*Fig. 2*), to broadly dysregulated pathways affecting at least two cell types (*Fig 3*). Remarkably, the affected pathways displayed substantial similarity across brain regions, with some showing consistent directional changes across cell types, while others exhibited complex patterns of distinct alterations in each cell type (*Supplementary Fig. 2; see Supplementary Table 12 for full list of overlapping pathways*).

**Figure 2.**
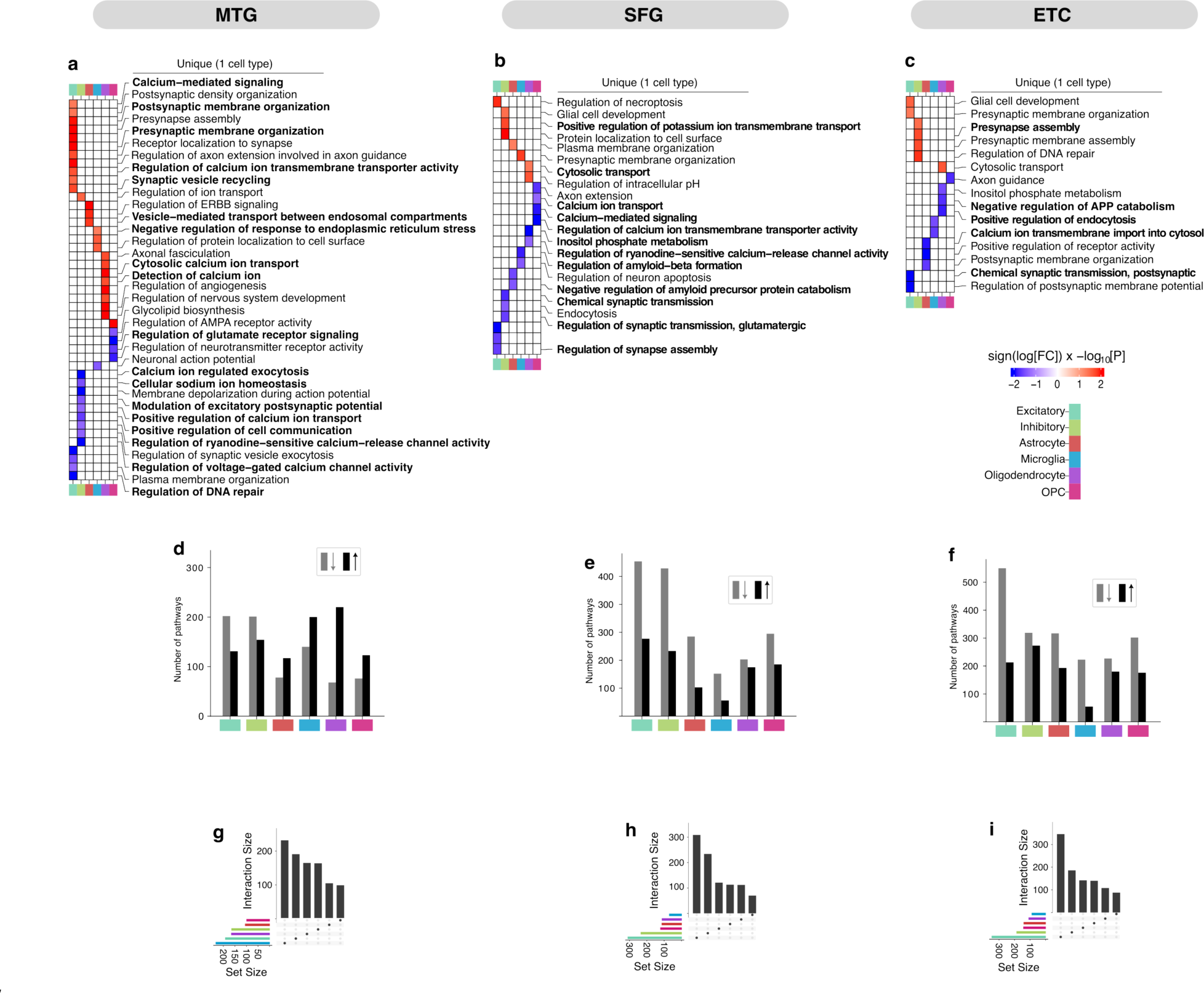
Cell-type-specific pathway perturbations in AD. (**a**—**c**). Heatmaps representing select Gene Ontology biological processes dysregulated in individual cell types (nominal P < 0.05). Each column represents data for a particular brain region. Unique alterations denote evidence of pathway alteration in a single cell type, with red indicating upregulation and blue signifying downregulation. Pathways discussed in the Article are highlighted in bold text. (**d—f**), Distribution of up- and down-regulated pathways across each cell type. Black indicates upregulation and grey indicates downregulation. (**g— i**), Upset plots displaying the distribution of uniquely perturbed pathways.

**Figure 3.**
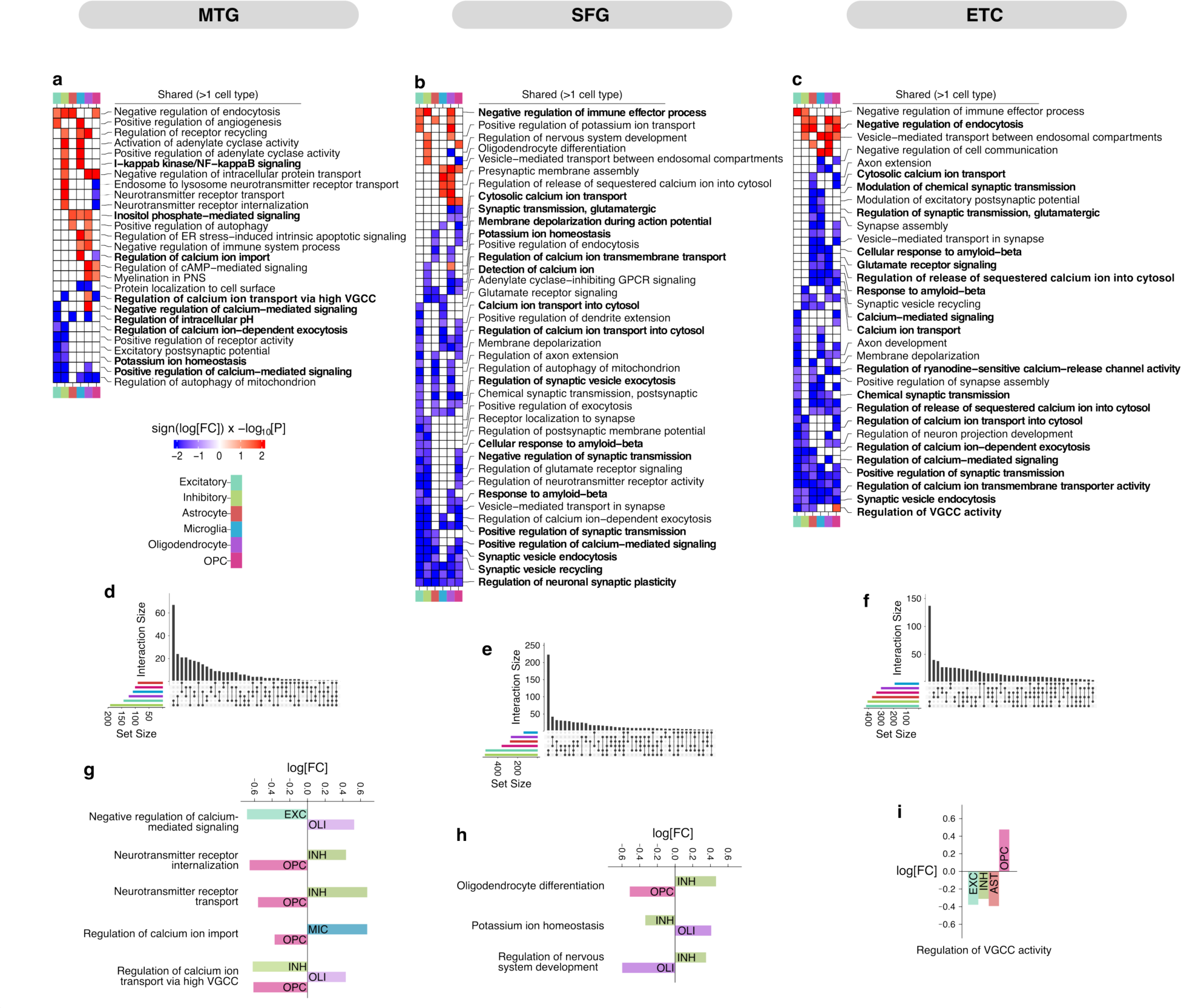
Broadly dysregulated processes in AD. (**a**—**c**). Heatmaps representing select Gene Ontology biological processes dysregulated in more than one cell type (nominal P < 0.05). Shared alterations are indicated as evidence of dysregulation in multiple cell types, with red representing upregulation and blue for downregulation. Pathways discussed in the Article are highlighted in bold text. (**d—f**), Upset plots displaying the broadly dysregulated pathways. (**g—i**), Selected pathways exhibiting different dysregulation patterns across cell types.

**Figure 4.**
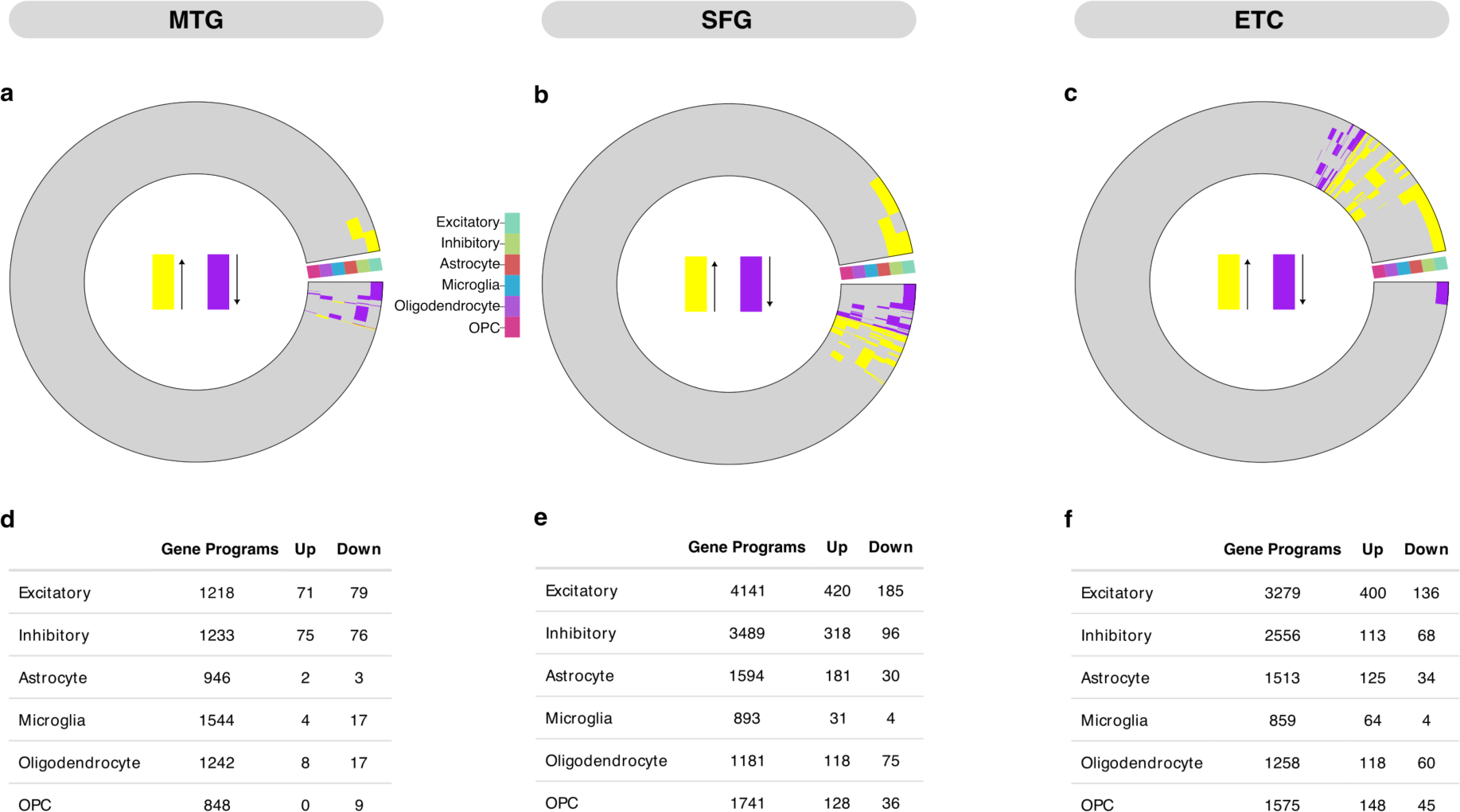
DEGs are underrepresented in perturbed gene programs. (a—c). Circular heatmaps illustrating cell-type specific dysregulation pattern of active genes amongst all perturbed pathways (unique and broad). Yellow and purple strips represent upregulated and downregulated genes respectively (false discovery rate (FDR) < 0.01 and abs(log2(fold change)>0.25), while grey regions represent non-DEGs, suggesting sparse presence of DEGs relative to total genes within the gene programs. (d—f). Table showing number of DEGs in the gene programs comprising perturbed pathways in each cell type.

Specifically, in the MTG, cell-type-specific perturbations were particularly evident in excitatory neurons, manifesting predominantly in dysregulation of synaptic-related processes, including upregulation of pre- and post-synaptic membrane organization, synaptic vesicle recycling, and Ca^2+^ - mediated synaptic signaling (*Fig. 2a*). Conversely, inhibitory neurons in the MTG showed a distinct pattern of downregulation in processes like excitatory postsynaptic potential, Ca^2+^ ion transport, and cell communication. Additionally, we observed cell-type-specific dysregulations in astrocytes (vesicle-mediated transport, *P=0.0149*), microglia (inflammatory response, *P=0.0003*), oligodendrocytes (nervous system development, *P=0.0049*), and OPCs (TORC1 signaling, *P=0.0001*), each demonstrating unique pathway alterations pertaining to their cellular functions. For instance, astrocytes exhibited upregulation of endosomal-related vesicle transport, while microglia showed alterations in protein localization and endoplasmic reticulum stress response (*Fig. 2a*). Oligodendrocytes were affected in nervous system development, and OPCs displayed perturbed glycolipid synthesis (*Fig. 2a*). However, a considerable number of processes were uniquely altered in specific cell types, highlighting the highly cell type-specific nature of pathway perturbations in the MTG (*Fig. 2g*).

Surprisingly, synaptic-related alterations were not exclusive to neurons; Ca^2+^ ion-dependent exocytosis was consistently downregulated across neuronal cells, while dysregulated neurotransmitter receptor transport and internalization were observed in OPCs and inhibitory neurons, among other broadly dysregulated processes (*Fig. 3a*). In addition, processes involved in the regulation of Ca^2+^ ions, voltage-gated Ca^2+^ channel activity and signaling, as well as myelination, exhibited distinct patterns of perturbation in excitatory, inhibitory, oligodendrocytes, and OPCs (Fig. 3g). Notably, 60% (n=333, P<0.05) and 57% of pathways are downregulated in excitatory and inhibitory neurons (n=355, P<0.05), respectively. In contrast 58-76% of pathways are upregulated in astrocytes (n=195, P<0.05), microglia (n=340, P<0.05), oligodendrocytes (n=287, P<0.05), and OPCs (n=199, P<0.05) (Fig. 2d). These findings together highlight the highly cell type-specific nature of pathway perturbations in the MTG, suggesting divergent mechanisms between neuronal and glial cells.

We next examined the pathway dysregulation patterns in the SFG and ETC. Similar to findings in the MTG, we observed a diverse set of AD-induced pathway alterations uniquely or broadly perturbed in the SFG and ETC (*Fig 2&3b,c*), further highlighting the heterogeneity of cellular responses to AD pathology. These include processes like synaptic transmission and membrane organization in neurons, amyloid beta formation and amyloid precursor protein (APP) catabolism in microglia, astrocytes, and oligodendrocytes, and axon maintenance processes regulated by OPCs. Interestingly, all cell types in both SFG and ETC exhibit a strong signature of repression, with 53-80% of pathways showing downregulation. This consistent alteration pattern across all cell types suggests a more pervasive disruption of molecular processes in these regions compared to the MTG. Moreover, a significant proportion of processes (53% in the SFG and 55% in the ETC) were perturbed either in neurons or a glial cell type, indicating a trend toward broader AD-driven disruption of molecular processes in the ETC and SFG (*Fig 3d—f*). Indeed, top differentially expressed pathways relevant to neuronal functions, such as Ca^2+^ -mediated signaling, Ca^2+^ ion transmembrane transporter activity, synaptic vesicle endocytosis, synaptic transmission, neuronal synaptic plasticity, and synaptic vesicle recycling, were consistently downregulated across all cell types in the SFG and ETC (*Fig 3b,c*). These pathways were predominantly downregulated in neurons, indicating that neuronal dysregulation dominates the AD-driven pathway alterations in the SFG and ETC. Concurrently, the observed changes in non-neuronal cell types appear to be closely associated with these neuronal perturbations.

To identify consistently perturbed processes across the three brain regions, we assessed the overlap of differentially perturbed pathways in a cell-type-specific manner (*Supplementary Fig. 2; see Supplementary Table 12 for full list of overlapping pathways*). We observed considerable concordance of altered cellular process across the three brain regions, with particularly notable overlaps between the SFG and ETC, likely reflective of reduced subject-specific variations. Interestingly, excitatory and inhibitory neurons showed the most pronounced concordance in disrupted processes (*Supplementary Fig. 2*a), with 31 pathways in excitatory neurons and 84 pathways in inhibitory neurons showing consistent dysregulations across the three regions. Alterations in inhibitory neurons include consistent downregulation of VGCC activity across all regions, and other key processes like Ca^2+^ -regulated signaling and exocytosis, potassium ion (K^+^) homeostasis, and neurotransmitter receptor maintenance. Excitatory neurons, on the other hand, consistently expressed disruptions in pathways related to mitochondrial autophagy and Ca^2+^-dependent exocytosis, underscoring the role of Ca^2+^ signaling in AD-associated cellular perturbations (*Supplementary Fig. 2*b). In contrast, glial cells exhibited less overlap in perturbations, with microglia cells, showing the least concordance. This suggests a broader spectrum of cellular disruptions and region-specific sensitivities to microglial dysregulation in AD. Among the affected processes were intracellular pH regulation, mitochondrial autophagy, and neurotransmitter receptor transport (*Supplementary Fig. 2*b). This variability among glial cells suggests a more complex and region-specific landscape of glial involvement in AD pathology.

Together, these findings elucidate the complex nature of cellular responses to AD, demonstrating that the cellular context in which AD manifests leads to markedly divergent molecular perturbations. The observed cell-type and region-specific perturbations highlight the complexity inherent in the regulatory landscape comprising the diverse molecular processes following AD pathogenesis.

### Unraveling AD-associated pathway alterations at systems level

We next ask whether molecular perturbations at the gene level are well represented in the biological processes dysregulated in AD. To accomplish this, we estimated cell-specific differences in gene expression between individuals with AD-pathology and healthy controls (**Methods**) and evaluated the enrichment of DEGs in perturbed processes. Surprisingly, our results reveal that only a small proportion of DEGs across all cell types (*Fig. 4a—c*) were associated with the gene programs (the curated set of genes comprising a biological process) implicated in the perturbed pathways earlier reported (*Fig. 2&3a—c)*. Notably, among the three brain regions, excitatory neurons in the ETC displayed the most, yet still limited representation, with only 16% of the 3,279 gene programs being DEGs (*Fig. 4d—f*). Neurons consistently showed the most significant degree of DEG representation with a combined 12% overlap in the MTG, 13% in the SFG, and 12% in the ETC. In contrast, astrocytes and microglia demonstrated markedly lower degree of DEG overlap, ranging from 0.5% to 13% across the three regions. These findings illustrate a sparse and varied representation of DEGs within perturbed processes, suggesting that relying solely on DEG analysis may not suffice to capture the full complexity of AD-related molecular changes.

Given the sparse and varied distribution of the DEGs within dysregulated processes, we sought to understand the extent to which DEGs influence the AD-associated perturbations at a systems-level. To achieve this, we examined the potential regulatory networks and overarching differences characterizing pathway disruption in AD across all cell types in these brain regions. We interrogated co-expression networks individually for each cell type in each brain region (*Fig 1, 5—8*), identifying groups of genes (gene modules) with high co-expression, suggesting potential co-regulatory mechanism or convergent biological functions. Specifically, network construction was confined to gene programs comprising earlier reported perturbed pathways (*Fig. 2,3)* enabling a fine-grained exploration of the molecular phenotypes governing the complex polygenic perturbations characteristic of AD. Traditional co-expression analysis methods developed for bulk transcriptomic data are not well suited to handle the inherent sparsity and noise in single-cell data (57,58). As a result, inferred networks are prone to spurious gene-gene correlations, thereby complicating the extraction of meaningful systems-level insights (59). To overcome these limitations, we estimate the gene-gene co-expression using hdWGCNA (31), a framework for co-expression network analysis tailored specifically for scRNA-seq data. hdWGCNA accounts for these considerations by aggregating highly similar cells into “metacells”, allowing for more accurate co-expression estimations, and facilitating the extraction of meaningful systems-level insights while preserving cellular diversity.

**Figure 5.**
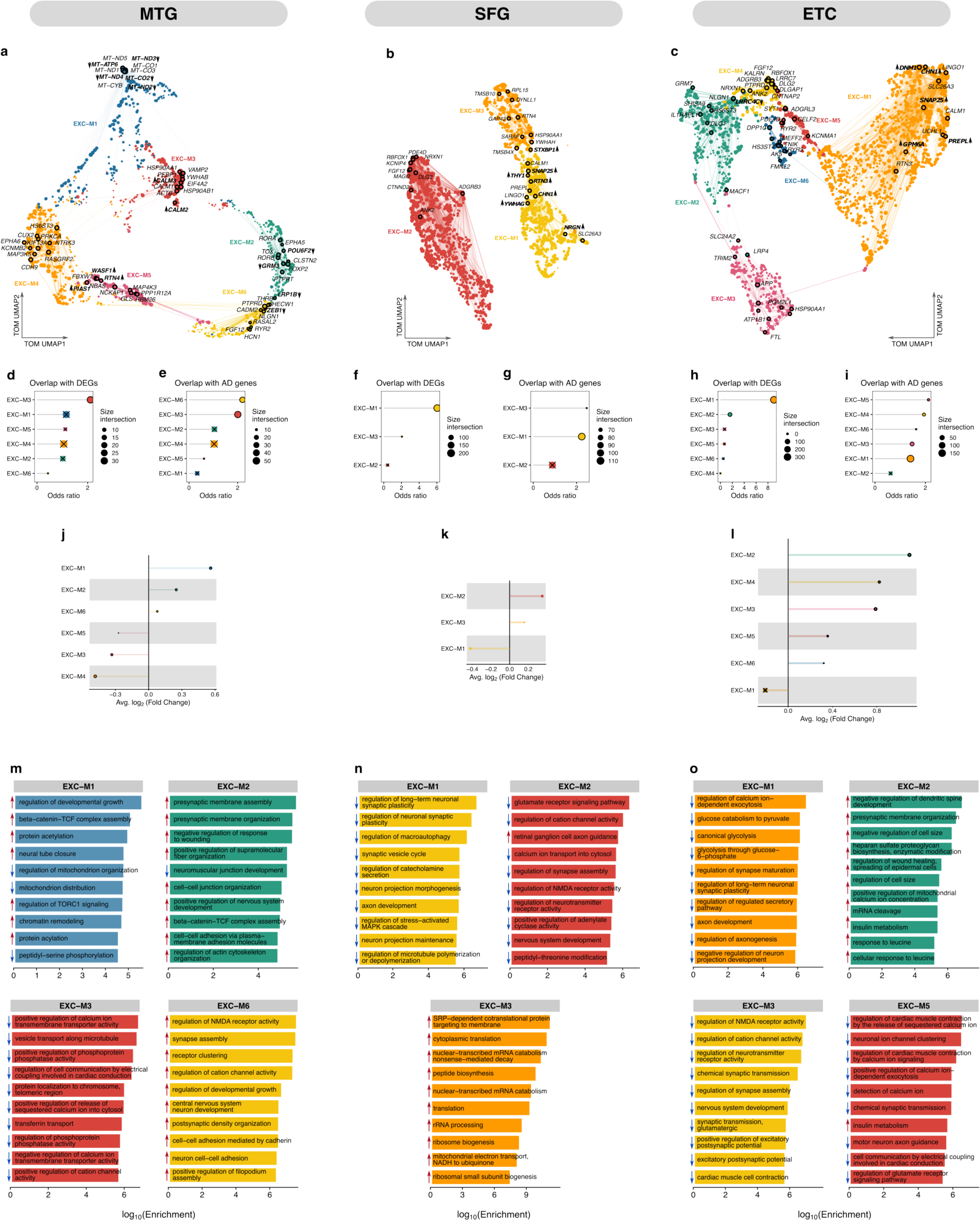
Disease-associated gene modules in excitatory neurons using co-expression networks derived from AD-dysregulated gene programs. (a—c). UMAP plot of the topological overlap matrix (TOM) illustrating neuronal co-expression networks constructed from genes programs comprising dysregulated pathways in excitatory neurons in the (a) MTG, (b) SFG, and (c) ETC. Nodes represent genes, color-coded by module membership, linked by edges depicting co-expression strength, with node size reflecting gene eigengene-based connectivity (**Methods**). Top hub genes are annotated within each module, with bold labels and directional arrows indicating hub-DEGs (hDEGs) as up- or down- regulated. Network visualization is simplified by edge downsampling for clarity. (d—i). Gene overlap analysis showing overlap of DEGs (d, f, h) and AD-associated genes (**Methods**) (e, g, i) with genes within co-expression modules, using Fisher’s exact test. An “X” indicates nonsignificant overlap (FDR > 0.05). (j—l). Lollipop plots representing the fold-change of differential expression for Module Eigengenes (DMEs), with the dot size corresponding to the number of genes in the respective module. An “X” overlays modules without statistically significant expression changes (FDR > 0.05). (m—o). Gene Ontology (GO) term enrichment within differentially expressed co-expression modules. Bar plots illustrate the log-scaled enrichment scores; blue arrows indicate downregulated, and red arrows indicate upregulated processes.

**Figure 6.**
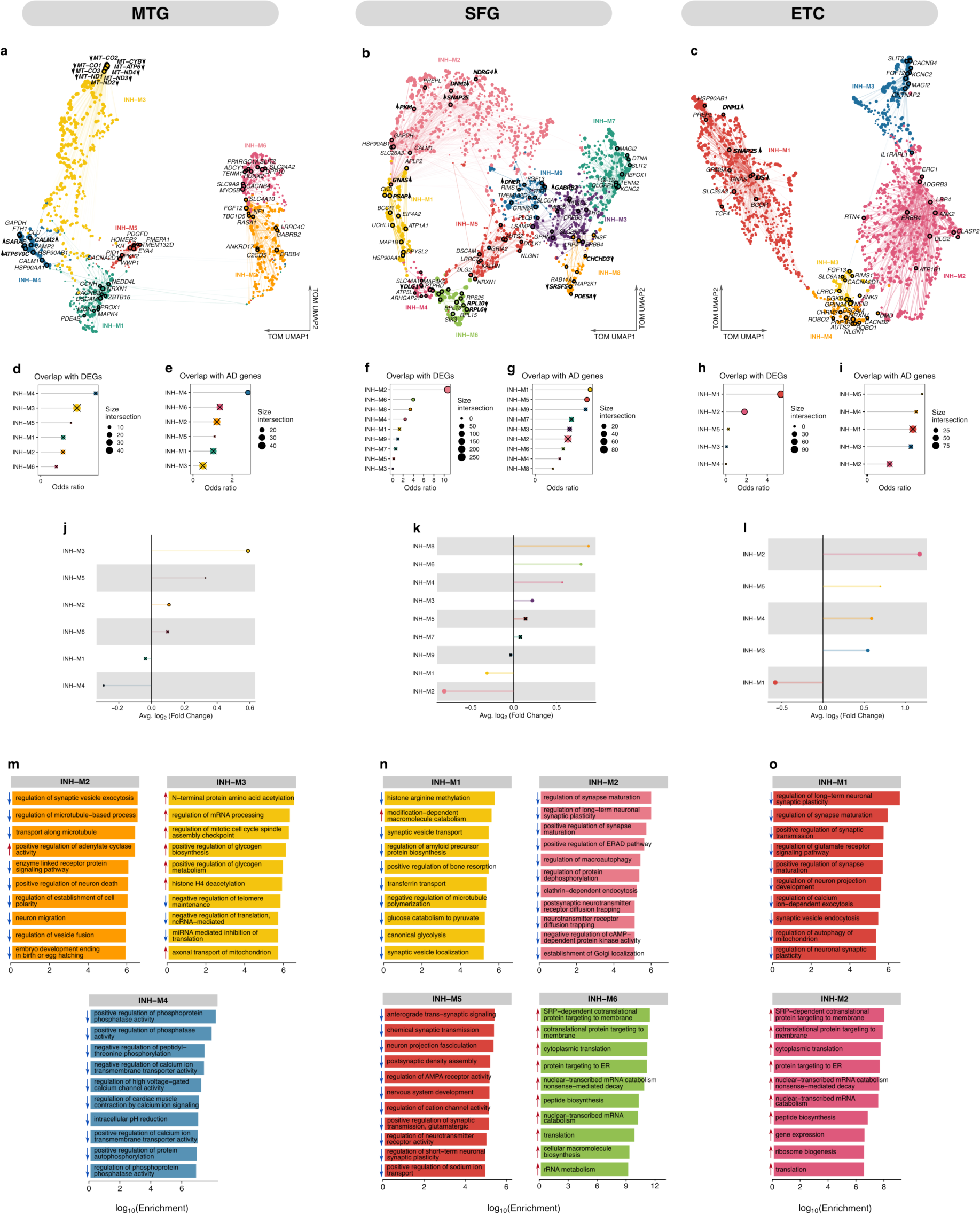
Disease-associated gene modules in inhibitory neurons using co-expression networks derived from AD-dysregulated gene programs. (a—c). UMAP plot of the TOM illustrating neuronal co-expression networks constructed from genes programs comprising dysregulated pathways in inhibitory neurons in the (a) MTG, (b) SFG, and (c) ETC. Nodes represent genes, color-coded by module membership, linked by edges depicting co-expression strength, with node size reflecting gene eigengene-based connectivity (see Methods). Top hub genes are annotated within each module, with bold labels and directional arrows indicating hDEGs as up- or down-regulated. Network visualization is simplified by edge downsampling for clarity. (d—i). Gene overlap analysis showing overlap of DEGs (d, f, h) and AD-associated genes (**Methods**) (e, g, i) with genes within co-expression modules, using Fisher’s exact test. An “X” indicates nonsignificant overlap (FDR > 0.05). (j—l). Lollipop plots representing the fold-change of DMEs, with the dot size corresponding to the number of genes in the respective module. An “X” overlays modules without statistically significant expression changes (FDR > 0.05). (m—o). GO term enrichment within differentially expressed co-expression modules. Bar plots illustrate the log-scaled enrichment scores; blue arrows indicate downregulated, and red arrows indicate upregulated processes.

**Figure 7.**
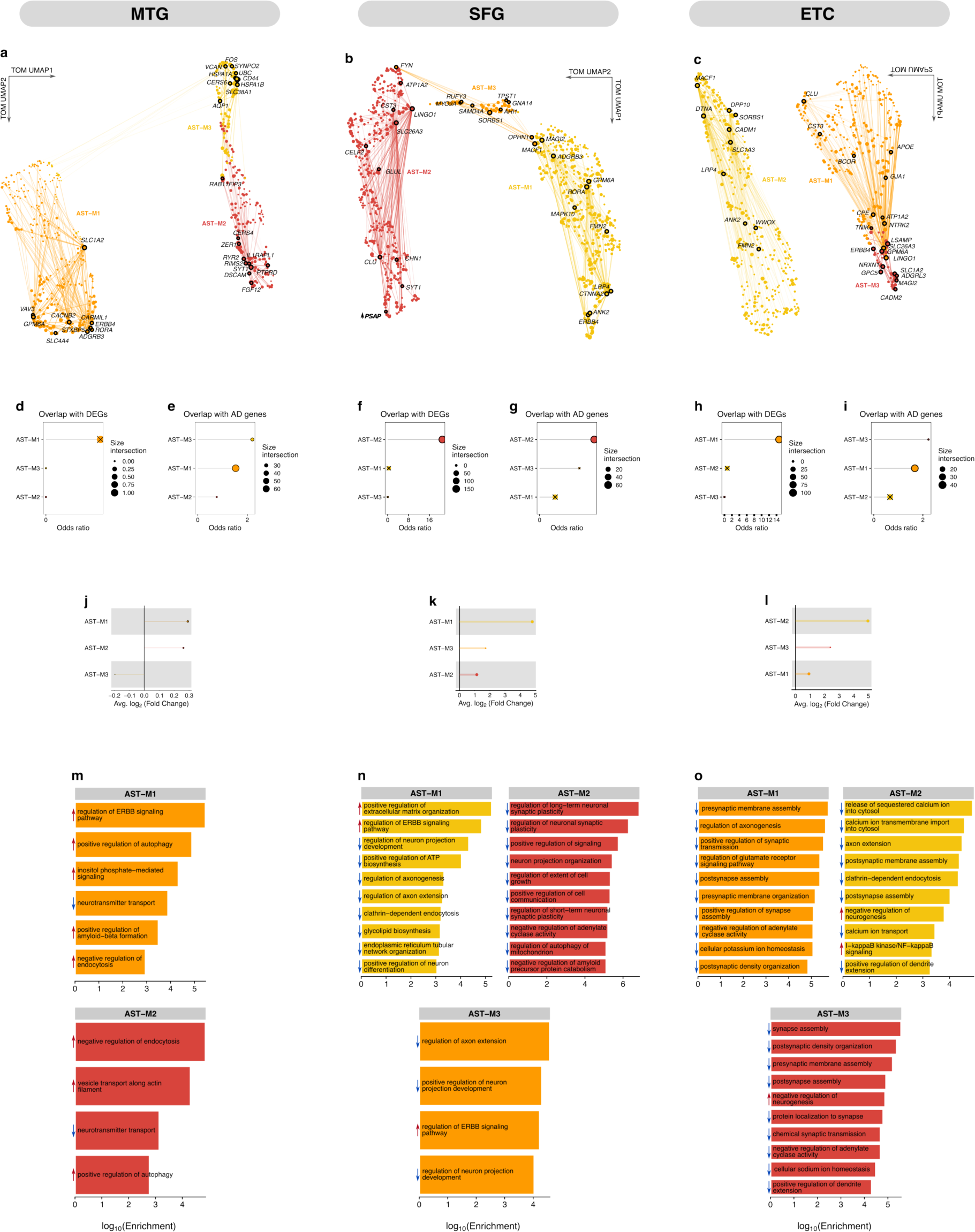
Disease-associated gene modules in astrocytes using co-expression networks derived from AD-dysregulated gene programs. (a—c). UMAP plot of the TOM illustrating glial co-expression networks constructed from genes programs comprising dysregulated pathways in astrocytes in the (a) MTG, (b) SFG, and (c) ETC. Nodes represent genes, color-coded by module membership, linked by edges depicting co-expression strength, with node size reflecting gene eigengene-based connectivity (see Methods). Top hub genes are annotated within each module, with bold labels and directional arrows indicating hDEGs as up- or down-regulated. Network visualization is simplified by edge downsampling for clarity. (d—i). Gene overlap analysis showing overlap of DEGs (d, f, h) and AD- associated genes (**Methods**) (e, g, i) with genes within co-expression modules, using Fisher’s exact test. An “X” indicates nonsignificant overlap (FDR > 0.05). (j—l). Lollipop plots representing the fold- change of DMEs, with the dot size corresponding to the number of genes in the respective module. An “X” overlays modules without statistically significant expression changes (FDR > 0.05). (m—o). GO term enrichment within differentially expressed co-expression modules. Bar plots illustrate the log- scaled enrichment scores; blue arrows indicate downregulated, and red arrows indicate upregulated processes.

**Figure 8.**
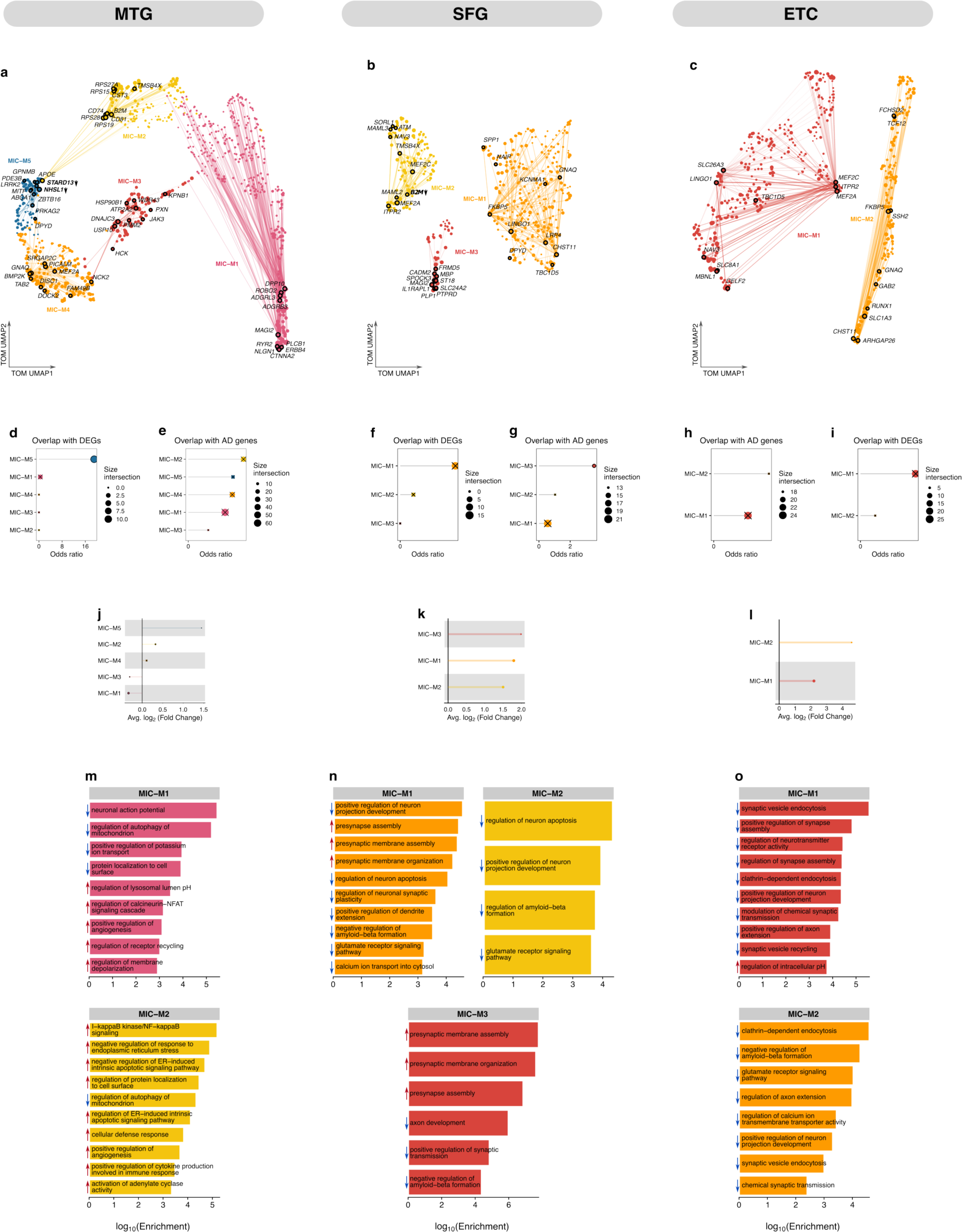
Disease-associated gene modules in microglia using co-expression networks derived from AD-dysregulated gene programs. (a—c). UMAP plot of the TOM illustrating glial co-expression networks constructed from genes programs comprising dysregulated pathways in microglia in the (a) MTG, (b) SFG, and (c) ETC. Nodes represent genes, color-coded by module membership, linked by edges depicting co-expression strength, with node size reflecting gene eigengene-based connectivity (**Methods**). Top hub genes are annotated within each module, with bold labels and directional arrows indicating hDEGs as up- or down-regulated. Network visualization is simplified by edge downsampling for clarity. (d—i). Gene overlap analysis showing overlap of DEGs (d, f, h) and AD-associated genes (Methods) (e, g, i) with genes within co-expression modules, using Fisher’s exact test. An “X” indicates nonsignificant overlap (FDR > 0.05). (j—l). GO term enrichment within differentially expressed co- expression modules. Bar plots illustrate the log-scaled enrichment scores; blue arrows indicate downregulated, and red arrows indicate upregulated processes. (m—o). Lollipop plots representing the fold-change of DMEs, with the dot size corresponding to the number of genes in the respective module. An “X” overlays modules without statistically significant expression changes (FDR > 0.05).

### Neuron-specific co-expression signatures in AD

Using hdWGCNA, we obtained a collection of gene-gene co-expression relationships across neuronal cells in each brain region (*Fig 5&6a—c*). Specifically, in the MTG, we identified 6 distinct excitatory co-expression modules (referred to as EXC-M1 to EXC-M6) (*Fig. 5a*) and inhibitory modules (INH- M1 to INH-M6) (*Fig. 6a*). Since functional insights within a co-expression network often stems from a selected set of nodes possessing high centrality (called hub genes), we reasoned that these hub genes are likely to play pivotal roles in cellular functions due to their extensive network interactions (31,60,61). The network plots highlight the top hub genes within each module, some of which exhibited differential expression (hDEGs, where h indicates that the gene is both a DEG and a hub gene; *see Supplementary Table 13 for full list of hub genes*).

Strikingly, in the MTG, our results show a concentration of downregulated hDEGs in EXC-M1 (*Fig. 5a*) and INH-M3 (*Fig. 6a*), primarily associated with cytosolic-localized RNA, such as MT-CO1, MT- ND3, and MT-ATP8. These genes encode essential subunits for oxidative phosphorylation in the electron transport chain, consistent with previous reports highlighting mitochondrial dysfunction, oxidative stress, and impaired cellular metabolism as key processes perturbed during AD pathogenesis (62). Similarly, we identified hDEGs in INH-M4 (*Fig. 6a*) and EXC-M3 (*Fig. 5a*), including members of the calmodulin gene family CALM2 and CALM3, recognized as regulators of intracellular Ca^2+^ signaling, with vital roles in synaptic processes. Previous studies have linked these hub genes to AD (Liu et al., 2020; Morabito et al., 2020; Wang et al., 2010), further substantiating the central role of Ca^2+^ signaling dysregulation in hippocampal AD pathogenesis, in accordance with the Ca^2+^ hypothesis of AD (63,66).

Furthermore, additional hDEGs were distributed across three excitatory neuron co-expression modules (EXC-M2, EXC-M5, EXC-M6) (*Fig. 5a*). Among these is the upregulated WASF1 in EXC-M5 (*Fig. 5a*), with a distinct regulatory role in actin assembly. Notably, downregulation of WASF1 has been linked to substantial reduction of amyloid levels within the hippocampus, indicating a negative feedback mechanism involving the APP intracellular domain—WASF1 pathway (67). The upregulated PIAS1 (EXC-M5), a known modulator of striatal transcription and DNA damage repair during SUMOylation, comprises critical parts of diverse cellular processes associated in neurodegenerative diseases like Huntington’s disease, Parkinson’s disease, and AD (68). Interestingly, PIAS1 overexpression was found to inhibit several AD marker genes such as NEUROD1, NEUN, MAPK2, GSAP, MAPT, and APP (69). Likewise, the downregulated ZEB1 expressed in EXC-M6 (*Fig. 5a*) underscores the role of transcriptional repression in regulating AD-associated correlations between accessible chromatin peaks and target genes (70). We next interrogated the distribution of DEGs and known AD-related genes using a comprehensive gene compendium from the Open Targets Platform (53), KEGG Alzheimer’s disease pathways (54), and Harmonizome (55) (*Fig. 5&6d—i*). Overlap analysis of modules in the MTG (**Methods**) revealed that, while up to 50 DEGs were distributed across excitatory and inhibitory co-expression modules, only EXC-M3 (*Fig. 5a*) exhibited significant enrichment for DEGs (*Fig. 5d*). Notably, three modules—EXC-M3, EXC-M6, and INH-M4—showed significant enrichment for AD-related genes (*Fig. 5&6d,e*). These results underscore the module-specific associations of DEGs and AD-related genes, suggesting intricate and dynamic transcriptional changes within co-expression modules and their potential relevance to AD pathogenesis. Additionally, we identified several AD-related hub genes distributed across excitatory and inhibitory co-expression modules (*Fig. 5&6a*). For instance, hub genes in EXC-M3 (*Fig. 5a*) included AD-associated genes HSP90AA1, HSP90AB1 which have been linked to protein misfolding, chaperoning, autophagy, apoptosis, and stress response—processes central to the dysregulation of protein integrity implicated in AD pathogenesis (71–73). Likewise, the presence of PPP1R12A in EXC-M5 (*Fig. 5a*) highlights its significance in the context of tau hyperphosphorylation and NFT formation, a hallmark of AD (74). Interestingly, EXC-M2 (*Fig. 5a*) expressed RORB, a classical marker of selectively susceptible excitatory neurons (20), while RYR2, expressed in EXC-M6 (*Fig. 5a*), regulates Ca^2+^ homeostasis and neuronal activity, which is central to normal cognitive function (75–77). INH-M4 (*Fig. 6a*) was enriched with key AD-related genes including GAPDH, CLU, and FTH1, involved in oxidative stress, amyloidogenesis, elevated cytotoxicity, and iron dysregulation, processes associated with AD progression. (78–81). Similarly, INH-M6 (*Fig. 6a*) contained DPP10, known to influence K^+^ channel activity and exhibit pronounced reactivity in the vicinity of NFTs and plaque-associated dystrophic neurites (82).

Next, we compared system-level differences in gene expression between AD and control groups using differential module eigengene (DME) analysis (**Methods**) (*Fig. 5&6 j—l*) (31). DME analysis of MTG derived neuronal modules revealed marked differences in the magnitude and direction of module expression from control to AD (*Fig. 5&6j*) (Wilcoxon rank-sum test Bonferroni-adjusted p <0.05; Supplementary Tables 9—11). These results suggest that AD-induced alterations in systems-level gene expression changes reflect either an enhancement of or decline in the functionality of co-regulated gene networks (83,84). Interestingly, all four down-regulated modules (*Fig. 5&6j*) (EXC-M3, EXC-M4, EXC-M5, INH-M4), exclusively comprised downregulated hDEGs (*Fig. 5&6a*). Conversely, upregulated modules (*Fig. 5&6j*) solely contained upregulated hDEGs (*Fig. 5&6a*), suggesting a pivotal role for hDEGs in perturbation of co-expression networks that characterize AD-related biological processes. Unsurprisingly, the downregulated excitatory module, EXC-M1, (*Fig. 5j*), enriched for mitochondrial-related hDEGs (*Fig. 5a*), was distinctively associated with differentially expressed pathways pivotal for numerous cellular processes and developmental functions (*Fig. 5m*). These include the regulation of development, assembly of the beta-catenin-TCF complex, protein acetylation, neural tube closure, mitochondrial organization and distribution, TORC1 signaling, chromatin alterations, protein acylation, and peptidyl-serine phosphorylation (*Fig. 5m*). Similarly, INH-M3 (*Fig. 5a*) was associated with genes that contribute to RNA processing, energy synthesis and metabolism, and protein stability (*Fig. 6m*) and was upregulated in AD (*Fig. 6j*). Moreover, other dysregulated modules (*Fig. 5&6i*) were found to be enriched for genes associated with a variety of biological processes crucial for normal neuronal functions, including synapse assembly (EXC-M2 & EXC-M6), vesicle transport (EXC-M3, EXC-M4, EXC- EXC-M7, EXC-M10, EXC-M13), Ca^2+^ transmembrane transport (EXC-M3, INH-M4), and synaptic vesicle exocytosis (INH-M2), which have been previously implicated in AD pathophysiology (*Fig. 5&6m*).

Co-expression analysis of neuronal cells in the SFG, resulted in a total of 12 excitatory and inhibitory modules (*Fig. 5b&6b*). Interestingly, EXC-M1 and M3 were significantly enriched in AD-associated genes and DEGs (*Fig. 5f,g*). Consistent with our observations in the MTG, AD-related hDEGs in the downregulated EXC-M1 (*Fig. 5k*), include SNAP25, NRGN, THY1, and RTN3, which are implicated in various processes central to AD pathophysiology, including synaptic neurotransmission, synaptic plasticity, synaptic signaling, immune response, neuron development, and endoplasmic reticulum (ER) morphology and function (*Fig. 5n*). Moreover, EXC-M2, though upregulated in AD (*Fig. 5k*), was associated with synaptic function, neuronal development, and signal transduction (*Fig. 5n*). A hub gene identified in EXC-M2, PDE4D, has been previously shown to result in abnormalities in the topological organization of functional brain networks (85). As a phosphodiesterase, PDE4D plays a pivotal role in regulating cAMP dynamics in neurons and glial cells (86), which ultimately influence memory formation and neuroinflammation (85,87). We noted GAP43, expressed in EXC-M3, whose elevated expression is recognized as a marker for tau and amyloid-driven pathologies. GAP43 also has a significant role in neural cell development, axonal sprouting, and regeneration (88–90). We also found enrichment of other AD-associated genes that have been prioritized as target genes in AD such as LINGO1 (EXC-M1) (91), NRGN (EXC-M1) (92), ADGRB3 (EXC-M2) (93), and RTN4 (EXC-M3) (94). This supports the notion that hub genes in these co-expression modules of the SFG are indicative markers of pathway dysregulation in excitatory cells in AD.

A total of 6 inhibitory modules were significantly enriched for DEGs or AD-related genes (*Fig. 6f,g*). In contrast to our observation in excitatory cells (*Fig. 5f,g*), none of these modules displayed simultaneous enrichment for both DEGs and AD-related genes (*Fig. 6f,g*). We observed a consistent pattern—either upregulation or downregulation—in the directionality of the hDEGs within their respective modules (INH-M1, INH-M2, INH-M4, INH-M6, INH-M8, and INH-M9) (*Fig 6b*). Surprisingly, and contrary to observations in the MTG (*Fig. 5&6a*), all hDEGs in both excitatory and inhibitory networks were counter-directional to the differentially expressed module eigengenes (DMEs) (*Fig. 5&6b*). This finding emphasizes the central role of hDEGs in the dysregulation of co-expression networks within AD-related pathways and suggests a robust region-specific association between hDEGs and module dysregulation. Notably, upregulated hDEGs such as SNAP25 and DNM1 in the INH-M2 (*Fig. 6b*), play critical roles in regulating synaptic vesicle fusion and recycling (95,96). Also, PKM, which is upregulated in INH-M1 (*Fig. 6b*) and is involved in glycolysis, is associated with aberration role in the regulation of metabolism and synaptic function in AD (*Fig. 6n*) (97–99). Moreover, the presence of upregulated PSAP in INH-M1 (*Fig. 6b*), underscores its role in lysosomal catabolism of glycosphingolipids (*Fig. 6n*) (100), and further highlights its significance in lysosomal dysfunction and neuronal survival in AD (101). We also observed enrichment of other key AD-related hub genes, particularly those regulating synaptic function in INH-M5 and INH-M4 (*Fig. 6b,n*). These include DLG1, a hDEG in INH-M4, DLG2, GRIA2, NLGN1, and NRXN1, in INH-M5 (102). Similar to the enrichment of RNA processing observed in EXC-M3, we detected a significant presence of ribosomal related genes in INH-M6, notably the downregulated hDEGs RPL6 and RPL10 (*Fig. 6b,n*). These genes are critical for protein synthesis and have been linked to regulation of metal ion homeostasis and cell death in AD (102–104).

Interestingly, we observed a recurring theme in the neuronal co-expression networks in the ETC (*Fig. 5&6c*). hDEGs identified in the SFG, including SNAP25, DNM1, CHN1, and DNM1 were also found to be hDEGs in EXC-M1 and INH-M1 in the ETC (*Fig. 5&6c*). In the same vein, all hDEGs across EXC-M1 and INH-M1 exhibited opposing directionality compared to the DMEs (*Fig. 5&6c,l*). Additionally, other AD-related hub genes were found to be shared across excitatory modules in both brain regions (*Fig. 5b,c*). These genes, including GRIN2A, NLGN1, NRXN1, and SLC6A1, play critical roles in synaptic formation, function, signaling, and plasticity. Notably, the heat shock protein HSP90AB1, essential for protein folding, was also identified as an AD-related hub gene shared among the excitatory modules. Similarly, we identified shared inhibitory hub genes with relevance to AD, such as CALM1, HSP90AA1, PDE4D, NRXN1, and RTN3. These genes assume particular significance in the perturbation of biological processes in AD due to their involvement in Ca^2+^ signaling, tau pathology, synaptic function, cAMP modulation, protein aggregation, and neurotransmission. Furthermore, analysis of DME patterns across the three brain regions revealed a unique relationship between gene co-expression modules and associated pathways (*Fig. 5&6j—o*). Remarkably, dysregulated modules exhibited a predominant enrichment for pathways perturbed in a specific direction. For instance, across excitatory and inhibitory modules in the ETC, the top enriched pathways were either consistently downregulated (EXC-M1, EXC-M3, and INH-M1), consistently upregulated (EXC-M2 and INH-M2), or predominantly downregulated (EXC-M5) (*Fig. 5&6o*). Interestingly, while certain DMEs displayed opposing dysregulation patterns relative to their corresponding enriched pathways, others demonstrated concordant dysregulation with enriched processes (*Fig. 5&6j—o*).

Taken together, these findings highlight the centrality of hDEGs and other AD-associated hub genes in orchestrating neuronal perturbations underlying the biological processes disrupted in AD. Hub genes identified within these networks shed light on the mechanisms of synaptic function, protein folding, and signaling that are significantly perturbed in neurons in AD. Furthermore, the alignment between dysregulated modules, enriched pathways, and hDEGs reinforces the notion of AD as a systems disease, characterized by tightly linked alterations in gene networks and their associated functional pathways.

### Glial-specific co-expression signatures in AD

To conceptualize the AD-driven systems-level perturbations in glial cells, we next probed the astrocyte (AST-M) and microglia (MIC-M) co-expression modules (*Fig. 7&8*). Contrary to the previously characterized neuronal co-expression patterns in the MTG (*Fig. 5&6d,e*), we observe that only one glial module, MIC-M5 (*Fig. 8a*), displayed significant enrichment for DEGs, and notably, was the only module containing hDEGs (*Fig. 7a,b,c,f,g,h*). This suggests a potentially limited role of DEGs in orchestrating systems-level differences in glial cells. While all astrocyte and microglia modules that displayed significant dysregulation in AD (*Fig. 7&8j*), contain AD-related hub genes (*Fig. 7&8a*), only AST-M3 and AST-M1 were predominantly enriched for AD-associated genes (*Fig. 7e*). Specifically, AST-M1, upregulated in AD (*Fig. 7i*), contained critical hub genes such as the glial high-affinity glutamate transporter, SLC1A2, a gene linked to altered glutamate homeostasis in AD and fundamental for preventing excitotoxicity in astrocytes and neurons (105,106); SLC4A4, a key regulator of neuronal pH homeostasis; and others including GPM6A, STXBP5, CACNB2, and ERBB4, which play roles in neurodevelopment, synaptic function and plasticity (107–111). Furthermore, AST-M2, which was upregulated in AD, contained hub genes linked to processes such as intracellular protein recycling (RAB11FIP3), immune response regulation, neuronal development, synaptic plasticity (IL1RAPL1, PTPRD), synaptic vesicle release (RIMS2, SYT1), and Ca^2+^ signaling (RYR2) (*Fig. 7a*). Conversely, the downregulated astrocyte module M3 (*Fig. 7j*) was enriched for stress-response genes including heat-shock genes (HSPA1A, HSPA1B), and genes critical for extracellular matrix organization and cellular adhesion (VCAN, CD44) (*Fig. 7a*).

Remarkably, hub genes in MIC-M2 included classical markers of DAM, such as APOE, B2M, CST3, and CD81, along with genes involved in RNA and ribosomal processing (RPS27A, RPS15, RPS19, and RPS28) (*Fig. 8a*). This supports the notion that system-level upregulation observed in MIC-M2 is linked to the dysregulated immune response and activation of phagocytic states in microglia. Indeed MIC-M2 displayed enrichment for processes related to microglial inflammatory activation, including pathways like I−kappaB kinase/NF−kappaB signaling, ER stress response regulation, and ER−induced apoptotic signaling, along with modulation of adenylate cyclase activity and mitochondria autophagy (*Fig. 8m*). Similarly, hub genes in MIC-M5 were primarily associated with lipid processing and immune response (*Fig. 8a*). These included genes such as GPNMB, LRRK2, MITF, ABCA1, STARD13, ZBTB16, and PRKAG2. Notably, GPNMB, a critical regulator of microglial activation and neuroinflammation, has been demonstrated to stimulate the production of pro-inflammatory cytokines, thus contributing to the inflammatory cascade observed in AD (112–115). Further reinforcing the activated state of microglia, MIC-M4 contained critical hub genes like PICALM, which governs clathrin-mediated endocytosis and is fundamental to Aβ clearance (116,117); DOKC2, a key regulator of Rho GTPase activation, which are essential components in immune cell trafficking and microglial mobility (118,119); and TAB2, a multi-functional adaptor protein involved in multiple cellular stress response pathways including TGF-beta and NF-kappaB signaling. Conversely, MIC-M3 and MIC-M1 were significantly downregulated (*Fig. 8j*) in AD and contained hub genes involved with protein folding, stress response, intracellular signaling, signal transduction, and synaptic function (*Fig. 8a,m*). Particularly, NLGN1 in MIC-M1, which is critical for the formation and maintenance of synapses, emphasizes the role of microglia in synaptic pruning and modulating neuronal connectivity. Additionally, the presence of RYR2 and PLCB1 in MIC-M1 suggests a crucial role for Ca^2+^ -mediated signaling pathways in modulating neuroinflammation and phagocytic activity of microglia.

Co-expression analysis of astrocyte and microglial gene programs in the SFG revealed 6 distinct modules (*Fig. 7&8a*). Strikingly, DME analysis revealed that all astrocyte and microglia modules are downregulated in AD (*Fig. 7&8k*), consistent with the predominant pattern of pathway downregulation observed in SFG (*Fig. 2e*).

This reinforces the link between gene co-expression networks and the orchestration of functional perturbations of biological processes in AD. Surprisingly, AST-M2 emerged as the only module exhibiting significant enrichment for AD-associated genes and DEGs (*Fig. 7&8f,g*), implying a critical role for AST-M2 in orchestrating astrocytic function in the context of AD. Specifically, AST-M2 contained several AD-associated hub genes with distinct functional relevance. For instance, SYT1, a key regulator of synaptic vesicle exocytosis and neurotransmitter release (120), and LINGO1 associated with the perturbation of neural growth and AD-associated myelination defects in AD (91), are hub genes in AST-M2. In addition, ATP1A2, a gene essential for astrocytic regulation of neuronal excitability via the maintenance of \ K^+^ and Na^+^ homeostasis (121), was also identified as a hub gene in this module. Other hub genes included GLUL, essential for astrocytic clearance of synaptic glutamate (122), and PSAP, an upregulated hDEG, implicated in dysregulation of lysosomal function and lipid metabolism in AD (101,123). Notably, classical markers of reactive disease associate astrocytes (DAA), CST3 and CLU, were also hub genes in AST-M2, known for their involvement in the clearance and accumulation of Aβ (26). Indeed, GO term enrichment revealed a robust array of biological processes governed by AST-M2, including maintenance of synaptic plasticity, signaling cascades, neuronal growth and repair, intercellular communication, and Aβ aggregation and clearance (*Fig. 7n*). This spectrum of functions effectively contextualizes the role of AST-M2 in the astrocyte-mediated maintenance of synaptic function and overall neuronal health, thus highlighting the integral role of neuron-glial crosstalk in the perturbation of the functional dynamics underpinning AD-related processes (124–126). We also found enrichment of genes associated with synaptic organization, cellular communication, energy metabolism, and development of neural structures in AST-M1 and AST-M3 (*Fig. 7b*). Indeed, hub genes in these modules play crucial roles in AD-associated process, including FYN in AST-M3, implicated in abnormal phosphorylation of tau protein and mediation of Aβ toxicity (127,128). Additionally, MAPK10, a hub gene in AST-M1, is essential for signaling pathways that regulate various cellular processes, including synaptic plasticity, neuronal survival, and apoptosis (129–131).

Consistent with our findings in the MTG, the microglial networks in the SFG also contained hub genes involved in a variety of processes relevant to both AD and microglial activation (*Fig. 8b*). Hub genes in MIC-M1 included SPP1, NAIP, LINGO1, LRP4, TBC1D5, reflecting the enrichment for immune response, synaptic maintenance, and overall neuronal function (*Fig. 8b,n*). Notably, SPP in MIC-M1 is integral for the regulation of phagocytic markers, thus playing a vital role in synaptic engulfment in the presence of Aβ (132). Also involved in the regulation of autophagy is TBC1D5, a hub gene in MIC- M1, functioning as a molecular switch for membrane trafficking between endosomal and autophagosomal pathways (133). MIC-M2 featured hub genes SORL1 and B2M, both implicated in a variety of AD-related processes such as trafficking of APP and resultant amyloidosis in AD (134–138). These genes further underscore the role of endolysosomal—autophagic network in regulating microglial activation (139). Additionally, MIC-M3 displayed enrichment for processes including synaptic assembly and axon development, consistent with the presence of hub genes MBP, PLP1, and PTPRD. This observation also underscores the notion that microglia, while often characterized primarily by their role in immune response in AD, also engage an array of processes vital for maintaining neuronal integrity, such as neural development, synaptic organization, myelin formation and maintenance.

Glial co-expression patterns in ETC are similar to those observed in the SFG (*Fig. 7&8c*). Remarkably, all astrocytic and microglial modules in the ETC exhibited downregulation in AD (*Fig. 7&8l*) and were mostly enriched for downregulated pathways (*Fig. 7&8o*), reinforcing the prevailing theme of pathway downregulation witnessed in the ETC (*Fig. 2e*). AST-M1 in the ETC exhibited considerable concordance with AST-M2 in the SFG, with hub genes CLU, CST3, and APOE, reflecting the DAA signature (*Fig. 7c*). However, no microglia module showed significant enrichment for hub genes signaling activated microglia state (*Fig. 8c*). AST-M1 remained the only module demonstrating significant enrichment for both DEGs and AD-related genes (*Fig. 7h,i*). Consistent with previous observation in the SFG, this highlights a pivotal role for the observed co-expression patterns in regulating astrocytic functions in the context of AD. Additional astrocyte hub genes in the ETC, including NRXN1, CADM1, MACF1, MAGI2, LRP4, GJA1 and ADGRL3, have been identified as markers of a reactive astrocyte state, implicating them in amyloidosis, regulation of neuroinflammation, cellular interactions (Dai et al., 2023). Notably, cellular adhesion hub genes, CADMI and NRXN1, have been previously noted as critical for maintaining the synaptic integrity and are hypothesized to contribute to excitotoxicity by impairing the function of reactive astrocytes in the regulation of extracellular ion balance, pH, and glutamate concentration (141–143). We also identified a compelling cross-regional consistency with the identification of shared hub genes—ANK2, ATP1A2, CLU, CST3, ERBB4, FMN2, GPM6A, LINGO1, LRP4, MACF1, MAGI2, SLC26A3, and SORBS1—between the ETC and SFG modules (*Fig. 7b,c*). Given the data for both regions were obtained from the same cohort, these hub genes emerge as potential brain-wide markers for astrocytic reactivity in AD. Likewise, shared microglia hub genes include CHST11, FKBP5, GNAQ, ITPR2, LINGO1, MEF2A, MEF2C, NAV3, and TBC1D5 (*Fig. 8b,c*), revealing complex interplay of functional involvements, including extracellular matrix modification, G-protein signaling, and intracellular Ca^2+^ regulation.

These results provide a robust systems-level perspective on the functional diversity within astrocyte and microglial modules in AD. We identified specific modules in these glial cell types that exhibit perturbations and are enriched for glial-specific processes and hub genes, yet notably did not prominently feature hDEGs. This suggests that the pathophysiological mechanisms in astrocyte and microglia may rely more on the dysregulation of gene networks and associated pathways rather than isolated gene perturbations. For instance, microglial modules, such as MIC-M2 and MIC-M4 in the MTG, primarily feature non-DEGs linked to DAM activation and microglial inflammatory responses. This is complimented by the functional downregulation observed in MIC-M3 and MIC-M1, which, despite a lack of enrichment for DEGs, feature genes crucial for protein folding, cellular stress response, and synaptic maintenance. Likewise, astrocyte modules, though not enriched with hub-DEGs, display a spectrum of AD-related alterations peculiar to astrocytic functions, from glutamate homeostasis to intracellular protein recycling and stress response. Together, our results offer a robust framework for appreciating the role of genes in glial alterations associated with AD, extending beyond differential gene expression profiles to the broader systems-level interplay of gene interactions underpinning AD pathogenesis.

### Conserved molecular drivers underlying pathway dysregulation

Our analysis reveals pronounced modular heterogeneity and extensive functional disruptions in neurons and glia across the brain regions. To further identify potential common drivers directing these pathway perturbations across regions, we examined recurrent hub genes within each cell type (Supplementary Table 14). Excitatory neurons showed substantial overlap of hub genes mostly participating in Ca^2+^ regulation, autophagy, proteostasis, cell-cell adhesion, neuronal cell death, and synapse regulation. Notably, most of these hub genes were non-DEGs in at least one brain region yet are AD-related and co-expressed in similar modules across all three regions. This further reinforces the notion that coordinated dysregulation of genes within a module, rather than changes in select individual genes, may promote pathway perturbations. Particularly, 6 hub genes—ACTB, CALM1, CALM2, GAPDH, HSP90AB1, and UCHL1—consistently belong to the same module in each brain region. Given their known roles in Ca^2+^ signaling, protein homeostasis, and neuronal apoptosis, these genes likely serve as region-wide orchestrators directing alterations in neuronal pathways fundamental for normal function. Inhibitory neurons demonstrated comparable overlap of non-differentially expressed hub genes participating in Ca^2+^-mediated signaling and synaptic transmission. As with excitatory neurons, the majority of these hub genes are AD-related and co-expressed in the same module, including CALM1, HSP90AA1, PDE4D, and NRXN1. Their function in regulating critical neuronal processes likely positions them as potential conserved mediators of pathway disruptions.

Among glial cells, microglia exhibited the highest hub gene overlap consisting of GNAQ, MAML3, MEF2A, MTHFD1L, and TGFBR1. These genes govern an array of processes critical for microglial activation, including inflammation, immune responses, and signaling cascades, potentially indicating conserved mechanisms underlying microglial reactivity across brain regions affected in AD. Likewise, the only overlapping astrocytic hub genes, ERBB4 and GPM6A, assume extensive roles in pathways related to neuroinflammation and synaptic dysfunction.

Overall, our analysis of recurrent hub genes points to potential conserved orchestrators of pathway disruptions across brain regions in AD. Experimental validation of these predictions remains vital to firmly establishing their functional significance. Nonetheless, our multi-region analysis provides a foundation to guide future investigations into common mechanisms directing AD pathogenesis.

## Discussion

Here, we leverage pathway activity and gene co-expression analyses to delineate the complex, systems-level alterations that characterize AD neuropathology. While scRNA-seq has been pivotal in revealing the molecular signatures of AD, much emphasis has been placed on differentially expressed genes without a comprehensive examination of the role and functional interconnectivity among these genes in biological processes across brain regions and cell types. This limitation largely renders associated studies insufficient for capturing the complexity of AD as a systems disease. Utilizing snRNA-seq data profiled from postmortem brain samples of the middle temporal gyrus, superior frontal gyrus, and entorhinal cortex, we reveal an intricate dynamics of perturbed gene networks underpinning the pathology in both neuronal and glial cell types.

The pathophysiological landscape of AD is distinctly marked by cellular and regional heterogeneity, as demonstrated in this study. In the MTG, for instance, AD-induced dysregulations in synaptic functions were significantly more prevalent in neurons compared to glial cells, corroborating previous findings that implicate synaptic dysfunction as a key pathological feature of AD (10). Additionally, our observations of unique pathway dysregulations in glial cells in the MTG contribute to the emerging discourse on the role of glial cells in mediating synaptic impairment in AD etiology (144–146). In the SFG and ETC, we detect a broad downregulation of molecular pathways across multiple cell types, suggesting a more advanced and pervasive pathological state. This is consistent with the known sequential propagation of AD-related pathology across different brain regions (147,148). Interestingly, Ca^2+^ signaling emerged as a shared hub of dysregulation but manifests variably among cell types and regions, underlining opportunities for cell type- and region-specific interventions. This is of considerable interest as Ca^2+^ homeostasis is critical for various cellular functions and its disruption has been considered central to AD pathogenesis (63). We argue that such cellular and regional specificity could not only serve as unique biomarkers for disease states but may also be exploited for targeted drug development.

A critical observation in our study is the limited distribution of DEGs among the gene programs comprising the perturbed pathways. This underscores the limitation and inadequacy of conventional pathway analyses or DEG-centric approaches in fully elucidating the complex systems-level alterations characteristic of AD. Thus, our work here expands upon traditional differential expression analyses to capture intricate interplay within gene co-expression networks. As a result, we delineate AD-related hub genes within enriched co-expression modules, implicating a range of biological processes from cellular metabolism to oxidative stress and Ca^2+^ homeostasis. Such an expansive approach broadens the spectrum of putative therapeutic targets and underscores the necessity for systems-level intervention strategies. Importantly, our results demonstrate that AD inflicts a broad spectrum of functional perturbations of gene co-expression across the three brain regions. This heterogeneity in modular responses provides compelling evidence that AD represents collective molecular perturbations, encompassing a spectrum of disruptions across neuronal and glial cells. Notably, we identify distinct patterns of hub-DEGs in specific modules, with a predominant distribution in both excitatory and inhibitory modules, but markedly less presence in glial modules. This pattern suggests that while DEGs have a substantial impact on neuronal cells in the context of AD, their influence on glial cells appears more limited. Given the propensity of co-expression networks to operate as integrated biological units, these findings lend support to the hypothesis that DEGs exert a disproportionately significant impact on neuronal dysfunction vis-à-vis the broader systems-level perturbations characteristic of AD.

Our study revealed a significant degree of functional heterogeneity among identified hDEGs. For instance, the upregulated hDEGs, WASF1 and PIAS1 are associated with actin assembly and DNA repair, respectively—mechanisms previously implicated in various neurodegenerative conditions, including AD (67,68). Additionally, the downregulated ZEB1 points to the role of epigenetic modifications, like accessible chromatin peaks, in AD pathology (70). We also identified certain modules particularly enriched for known AD-related genes, highlighting module-specific correlations with AD-driven pathway alterations. Hub genes in these enriched modules, including HSP90AA1 and HSP90AB1, GAPDH, CLU, and FTH1, implies a complexity that may signify both causative and reactive changes in AD pathogenesis. Moreso, our analysis revealed a prominent theme of mitochondrial dysfunction, underscored by the downregulation of hDEGs such as MT-CO1, MT-ND3, and MT-ATP8 in neuronal modules. The presence of these hDEGs lends compelling credence to the hypothesis that aberrations in mitochondrial dysfunction, cellular metabolism, and oxidative stress are key features of the AD pathological cascade (62). We also observed key hDEGs belonging to the calmodulin gene family (CALM2 and CALM3) in neuronal modules. Given the well-established role of these genes in regulating intracellular Ca^2+^ signaling, this observation adds a new perspective to the Ca^2+^ hypothesis of AD and is consistent with earlier works implicating them in disrupted Ca^2+^ signaling (13,63–66).

Differential module eigengene analysis further reinforced the notion that AD-associated perturbations result in both upregulation and downregulation of gene modules, consequently affecting a range of cellular processes. This further illuminates the collective behavior of genes within each module, emphasizing either an enhancement or decline of the functional output of co-regulated modules in AD. For instance, in the MTG, the downregulated neuronal modules exclusively comprised downregulated hDEGs and vice versa, implicating these genes in the system-level disruptions of cellular processes, which are essential for normal neuronal functions. This exclusive alignment underscores a strong functional coherence within these modules, suggesting that these hDEGs could be critical regulators in the onset and progression of AD, likely indicating a coordinated modular response to AD. Remarkably, our findings show intriguing patterns of interregional consistency and complexity. Across all brain regions, dysregulated modules exhibited a predominant enrichment for pathways perturbed in a specific direction. Interestingly, while certain DMEs displayed opposing dysregulation patterns relative to their corresponding enriched pathways, others demonstrated concordant dysregulation with enriched processes. Moreover, we observe region-specific counter-directionality of hDEGs in relation to the DMEs in the SFG versus MTG.

Our analysis of glial co-expression signatures across the brain regions elucidates the complex and dynamic roles of astrocytes and microglia in AD. We observed that only a single microglial or astrocyte module in each brain region showed significant enrichment for DEGs and reason that these modules represent critical functional drivers of pathway dysregulation. Consistent with this, we observed a significantly reduced number of hDEGs across all glial modules, pointing towards a potentially diminished role of DEGs in orchestrating glial-associated systems-level differences in AD. Contrary to extant narratives that largely assign a neuroinflammatory role to glial cells, our data unveil robust enrichment for AD-related genes involved in a range of biological processes, from synaptic pruning and stress response to glutamate homeostasis and Ca^2+^ signaling. This suggests that alterations in modular gene expression contribute significantly to the pervasive involvement of glial cells in AD. Specifically in microglia, we noted the critical role of modules governing dysregulated immune responses, phagocytic activities, and synaptic function. Such findings underscore the multi- functionality of microglia in AD, highlighting their involvement in preserving neuronal integrity through synaptic maintenance, myelin formation, and other mechanisms. Additionally, our findings reveal the critical role of disease-associated glial states in AD pathology. We observed that hub genes in the AD-enriched glial modules were fundamentally associated with reactive astrocyte and microglia states, indicating that glial cells assume activated states due to the complex systems-level interactions among these genes.

Cross-regional analysis between the MTG, SFG, and ETC, reinforced the theme of overall downregulation of both astrocytic and microglial modules in AD, indicating a prevailing trend of functional repression in these glial cells. These observations collectively strengthen the notion of AD as a systems disease, characterized by tightly linked alterations in gene networks and their associated functional pathways. We also notably identified shared hub genes across these brain regions, with more prominent overlap in neurons. These conserved hubs likely orchestrate directing modular dysregulation and pathway perturbations linked to critical neuronal processes like Ca^2+^ signaling, proteostasis, inflammation, and synaptic function. Though not always differentially expressed themselves, their coordinated behavior within modules may underpin consistent pathway disruptions in AD. Glial cells express more limited overlap, but shared genes govern diverse glial activation-related processes, potentially serving as brain-wide markers for astrocytic or microglial reactivity for disease diagnostics or targeted therapeutic interventions. Nevertheless, experimental validation remains essential to confirm the role of these putative hub genes as conserved, causal drivers of AD pathogenesis. In summary, integrated analyses of cell type-specific co-expression modules across multiple affected brain regions hold significant potential for elucidating key network regulators and pathways that may offer new therapeutic targets for AD.

## Conclusions

Our study provides a comprehensive systems-level analysis of the pathway perturbations associated with AD across multiple brain regions and cell types. Leveraging snRNA-seq data, we integrate pathway activity analysis with WGCNA, revealing profound heterogeneity in the dysregulation of biological processes in neurons and glia. Synaptic dysfunction and dysregulated Ca^2+^ signaling emerging as convergent axes of pathogenesis. Surprisingly, we observe limited overlap between DEGs and disrupted gene programs, suggesting DEGs alone do not adequately represent the collective modular alterations driving AD pathology. Indeed, we demonstrate that DEGs have a more pronounced role in driving modular dysregulation in neurons compared to glial cells. We also identified conserved hub genes across modules and brain regions which offers potential brain-wide cell-type-specific therapeutic targets and biomarkers. Overall, these findings underscore the necessity of integrated, systems-oriented models to fully capture the complexity of molecular interactions underlying AD and other polygenic systems neurodegenerative disorders.

## List of abbreviations

AD: Alzheimer’s Disease
Aβ: Amyloid-beta
NFTs: Neurofibrillary Tangles
scRNA-seq: Single-cell RNA-sequencing
snRNA-seq: Single-nucleus RNA-sequencing
DEG: Differential Gene Expression
DAM: Disease-Associated Microglia
Ca^2+^: Calcium
hub-DEGs: hub DEGs
MTG: Middle Temporal Gyrus
SFG: Superior Frontal Gyrus
ETC: Entorhinal Cortex
ADNC: AD Neuropathologic Change
ACT: Adult Changes in Thought
ADRC: Alzheimer’s Disease Research Center
NDBB: Neurodegenerative Disease Brain Bank
SEA-AD: Seattle Alzheimer’s Disease
GSVA: Gene Set Variation Analysis
WGCNA: Weighted Gene Co-expression Analysis
hdWGCNA: High Dimensional Weighted Gene Co-expression Analysis
ME: Module Eigengenes
DME: Differential Module Eigengenes
VGCC: Voltage Gated Calcium Channel

## Declarations

### Data availability

Processed snRNA-seq data and metadata from Leng et al. (20) are available for download at the Synapse portal ((42), under ID syn21788402), under controlled use conditions. Processed data from Gabitto et al. (35) can be freely downloaded without registration from the Seattle Alzheimer’s Disease (SEA-AD) Brain Cell portal (45) and a public AWS bucket (43).

### Code availability

The codes used for the analyses in this study, along with detailed instructions for reproducing the results presented here will be available on GitHub (https://github.com/TemiLeke/systematic_ad_analysis) upon publication.

### Authors’ Contributions

All authors read and approved the final manuscript. T.A. and G.U. conceived and presented the idea.

T.A. and S.I.I processed the data. T.A. conducted the analysis and wrote the initial draft of the manuscript, with input from S.I.I and G.U. S.I.I. assisted with methodology and analysis. G.U. supervised the study. All authors discussed and interpreted the results.

## Supplementary Information

### Additional File 1

**Supplementary Table 1.** Sample metadata for all 20 donors, including post-mortem neuropathological assessments, clinical evaluations, and pathological grouping. **Supplementary Tables 2—4.** The table of p-values and log fold changes for all genes included in the differential analysis test across all brain regions (MTG 2; STG 3; ETC 4) and cell types. Supplementary Table 5. Table of pathway renaming conventions. **Supplementary Tables 6—8.** Comprehensive documentation of the pathway analysis results for each brain region (MTG 6; SFG 7; ETC 8) with detailed statistical results (coefficient estimates and p-values) for the prioritized candidate pathways identified across major cell types.

**Supplementary Tables 9—11.** Results from DME analysis for each brain region (MTG 9; SFG 10; ETC 11) across all cell types. **Supplementary Table 12.** List of overlapping dysregulated pathways along with corresponding statistics for each cell type. **Supplementary Table 13.** List of hub genes and hDEGs in each module for all cell types across brain regions. Overlapping hub genes are presented in **Supplementary Table 14.**

### Additional File 2

**Supplementary** Fig. 1. Principal Component Analysis (PCA) of aggregated pseudoreplicates highlighted with relevant metadata. **Supplementary Fig. 2**. Overlapping dysregulated biological processes across brain regions. **Supplementary Figs. 3—5**. Soft power thresholds demonstrating a fit to the scale-free topology model across all cell types in each brain region (MTG 3; SFG 4; ETC 5). **Supplementary Fig. 6—8**. Dendrograms showing the different co-expression modules resulting from the network analysis across cell types in each brain region (MTG 6; SFG 7; ETC 8).

---

**Supplementary Fig. 1.**
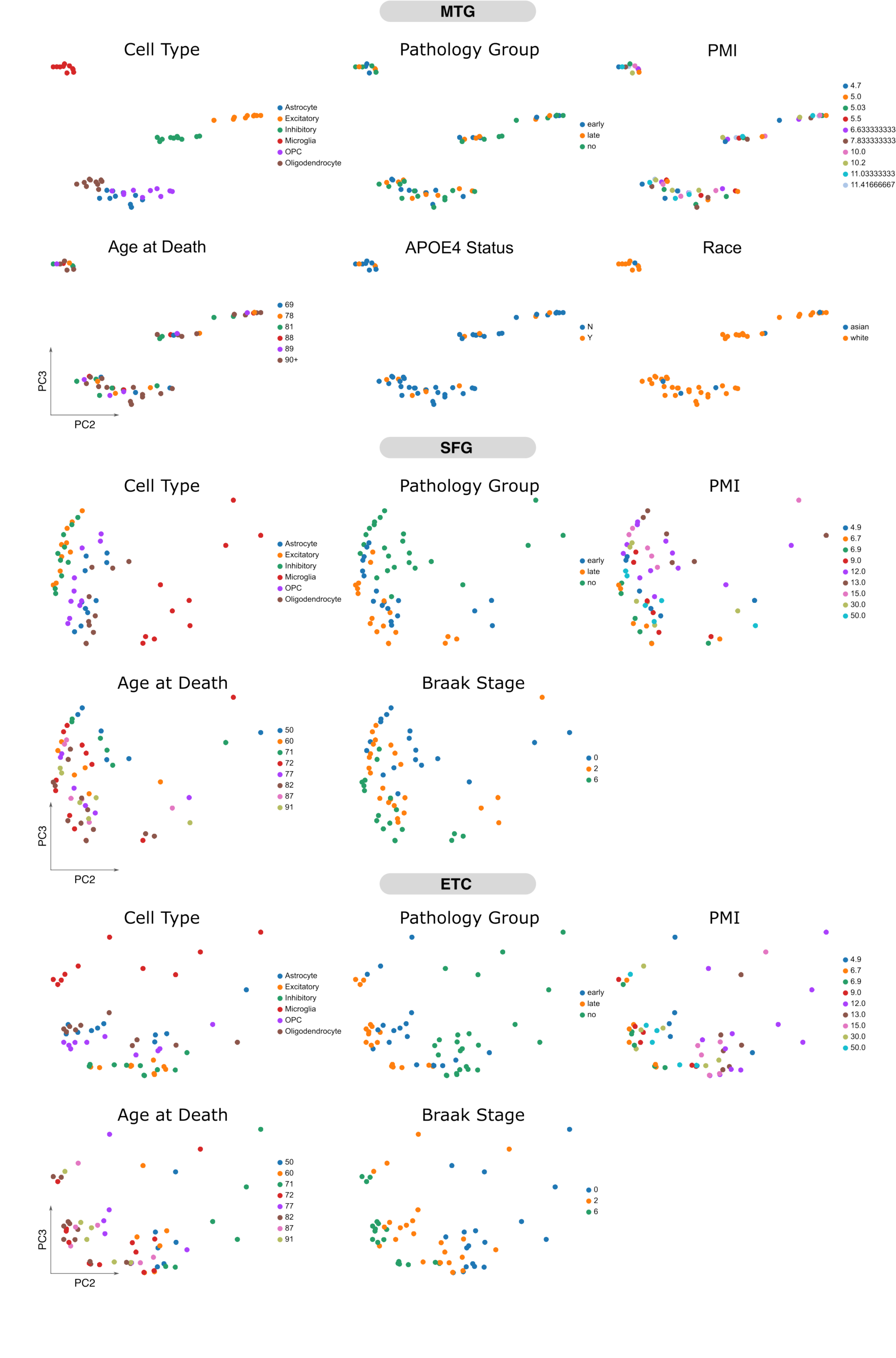

**Supplementary Fig. 2.**
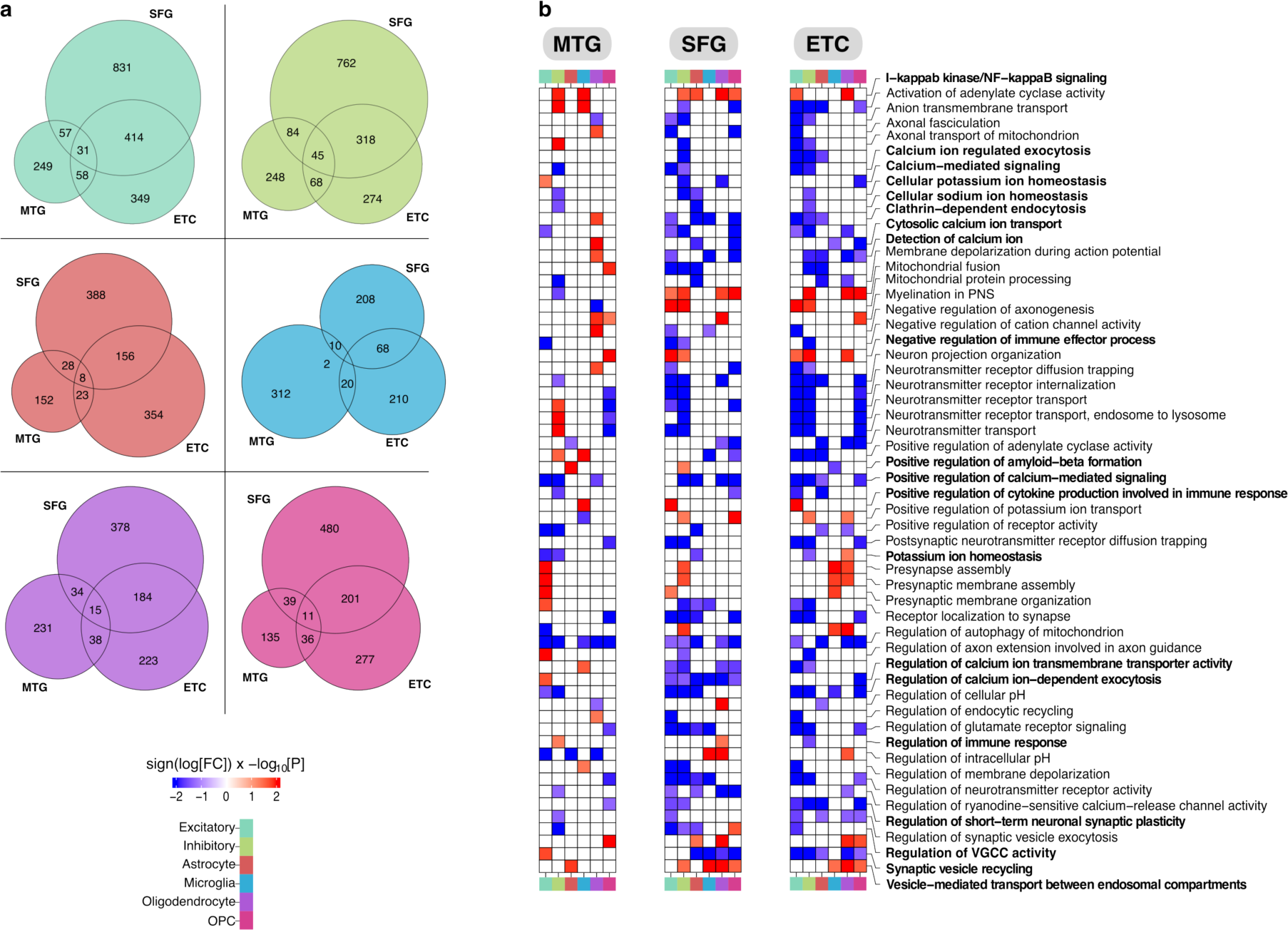

**Supplementary Fig. 3.**
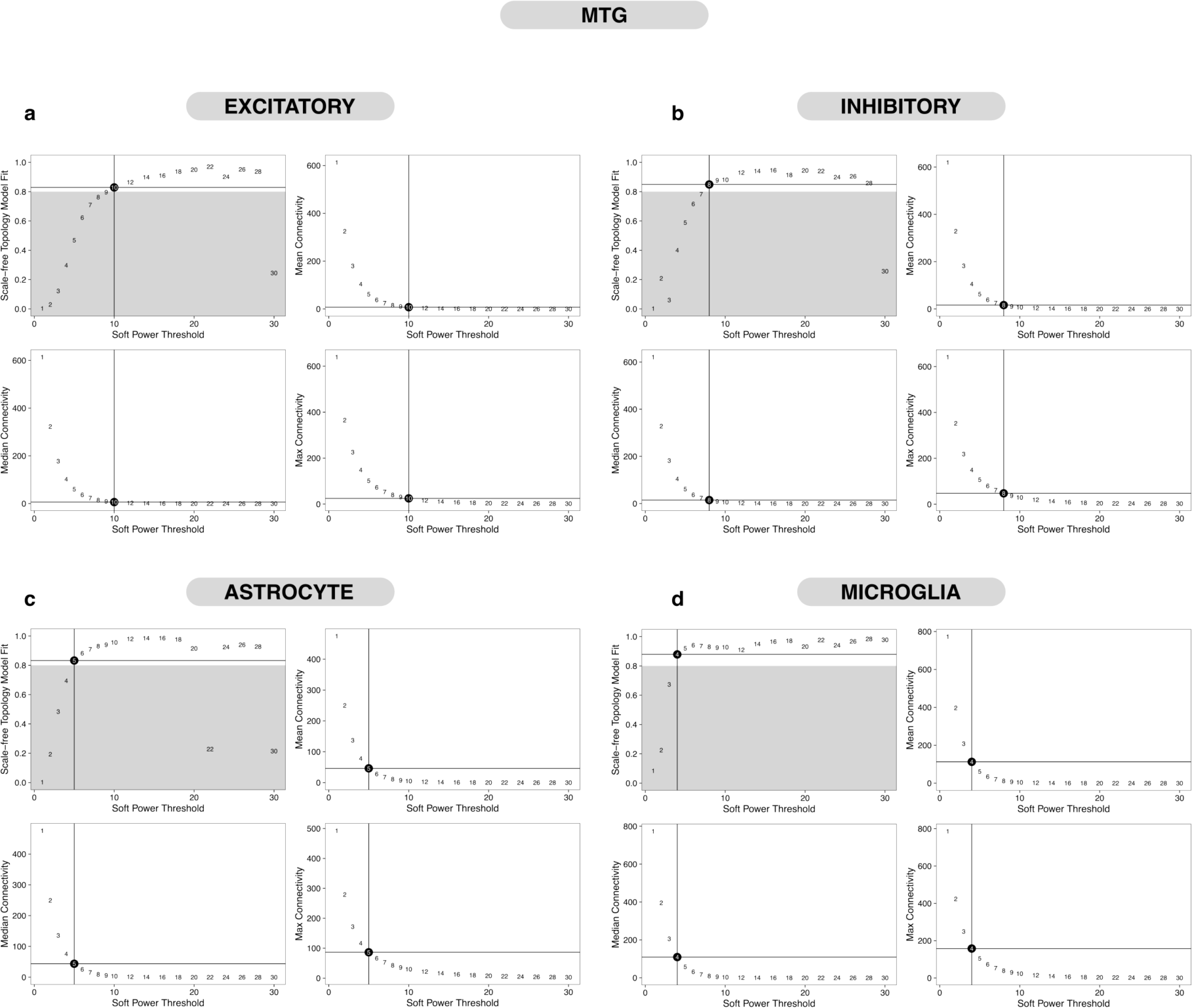

**Supplementary Fig. 4.**
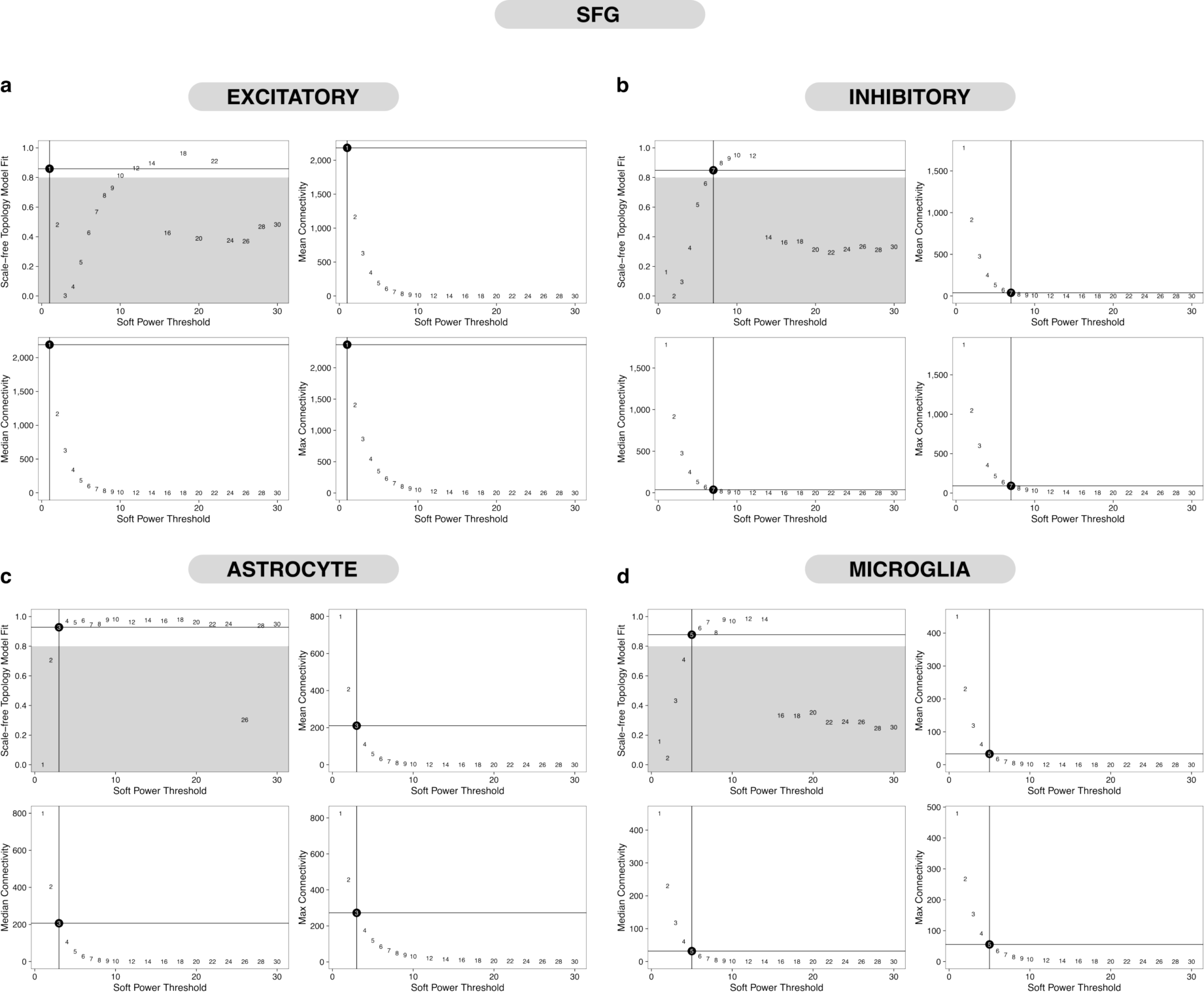

**Supplementary Fig. 5.**
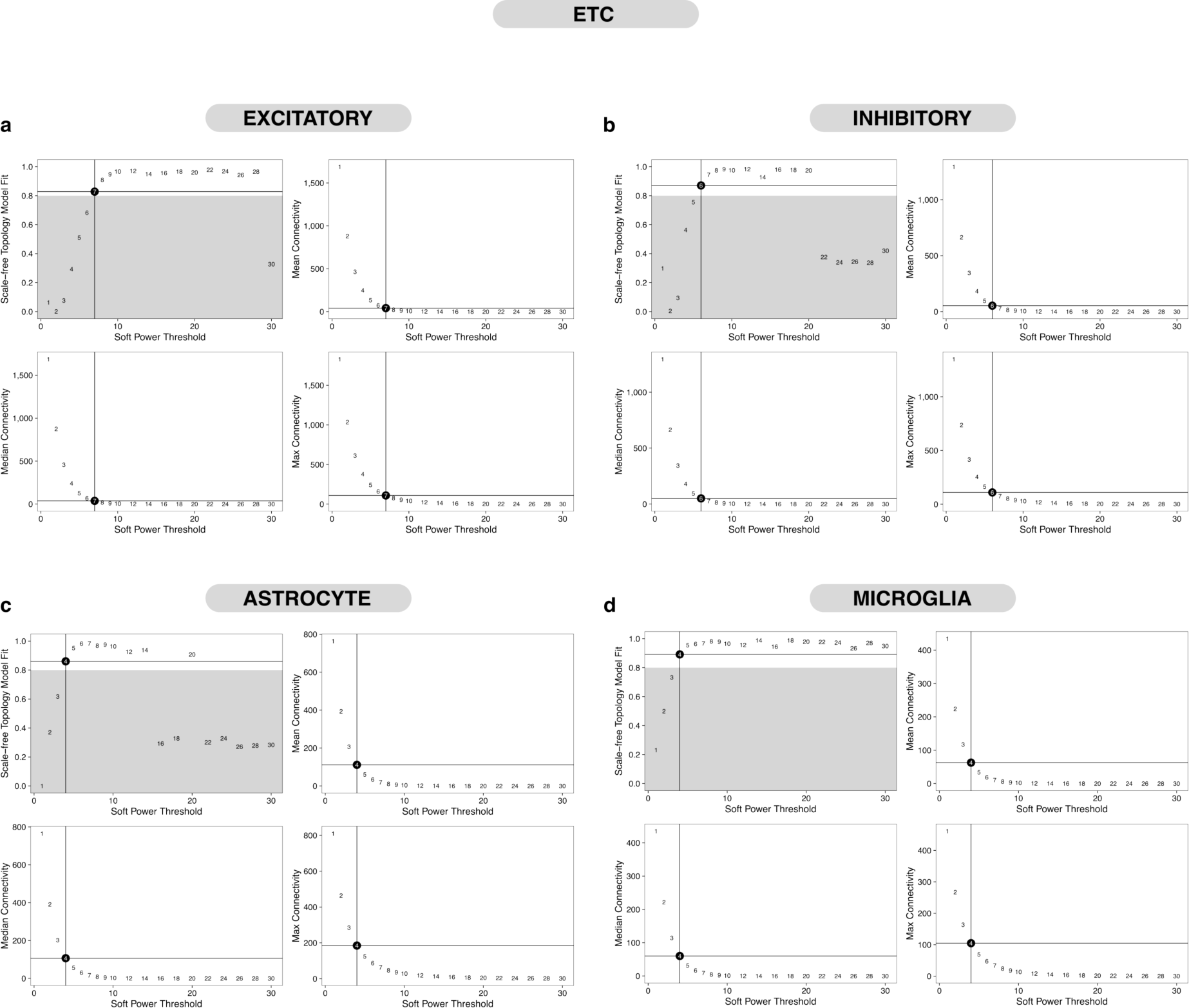

**Supplementary Fig. 6.**
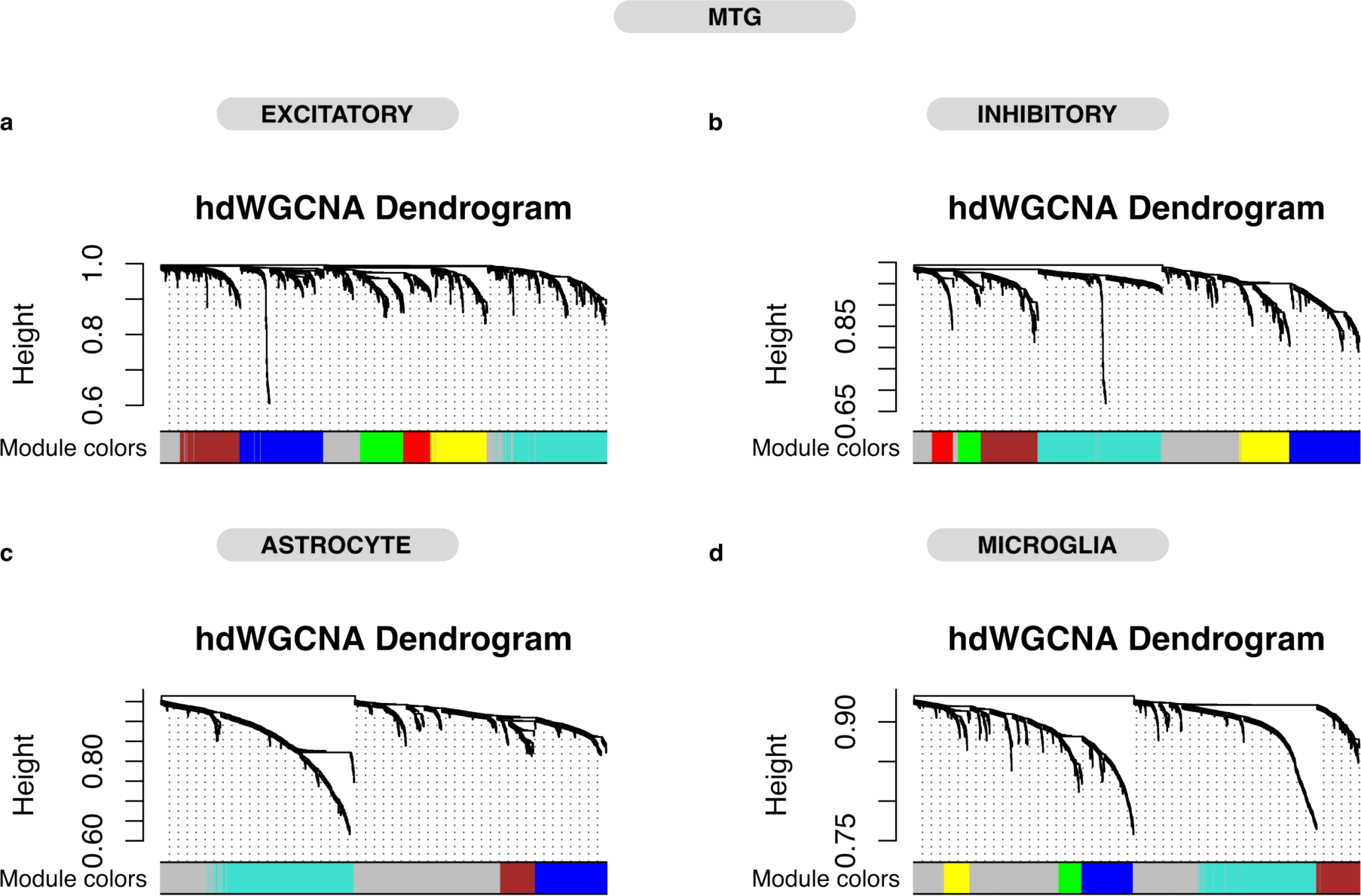

**Supplementary Fig. 7.**
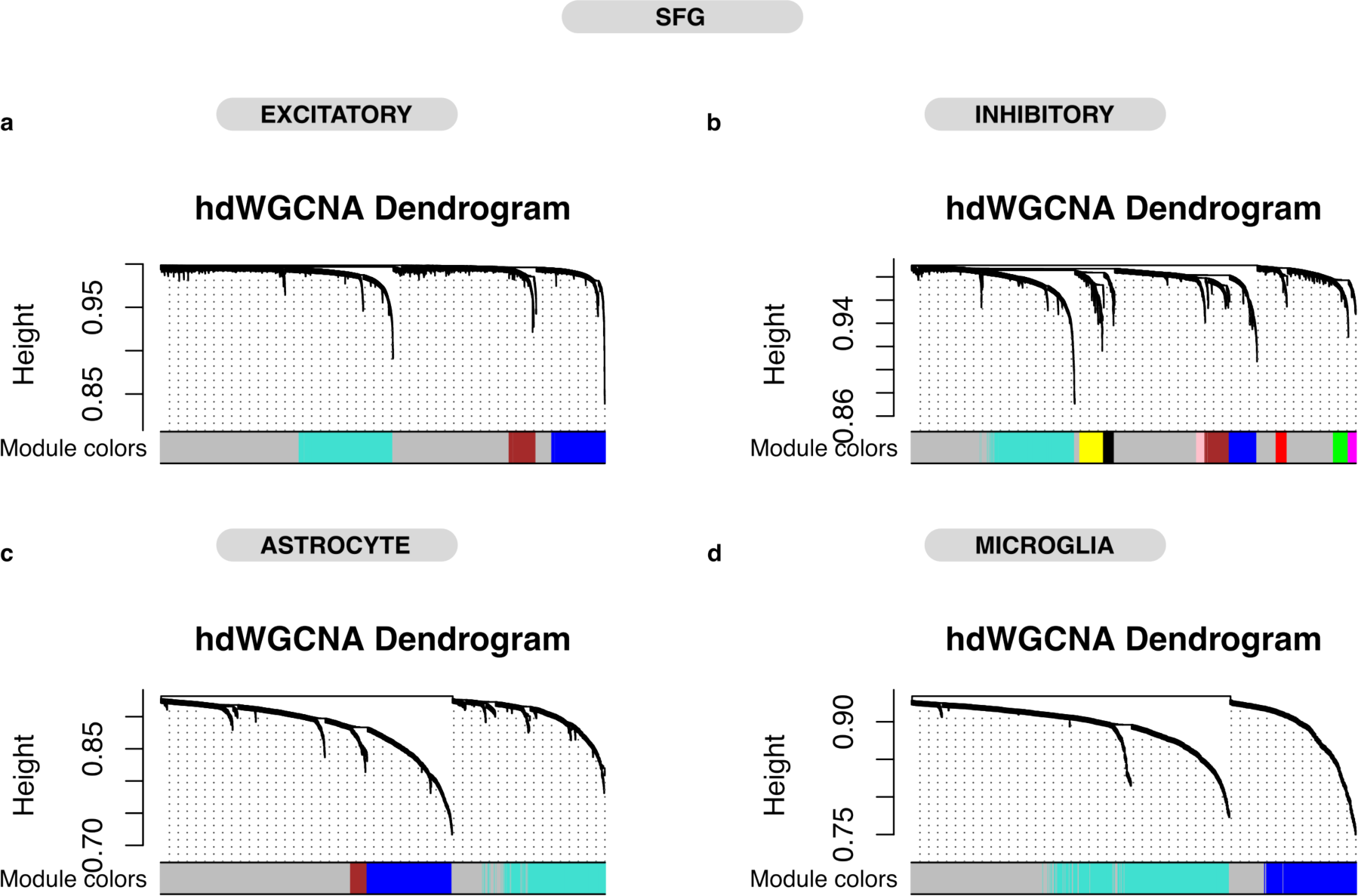

**Supplementary Fig. 8.**
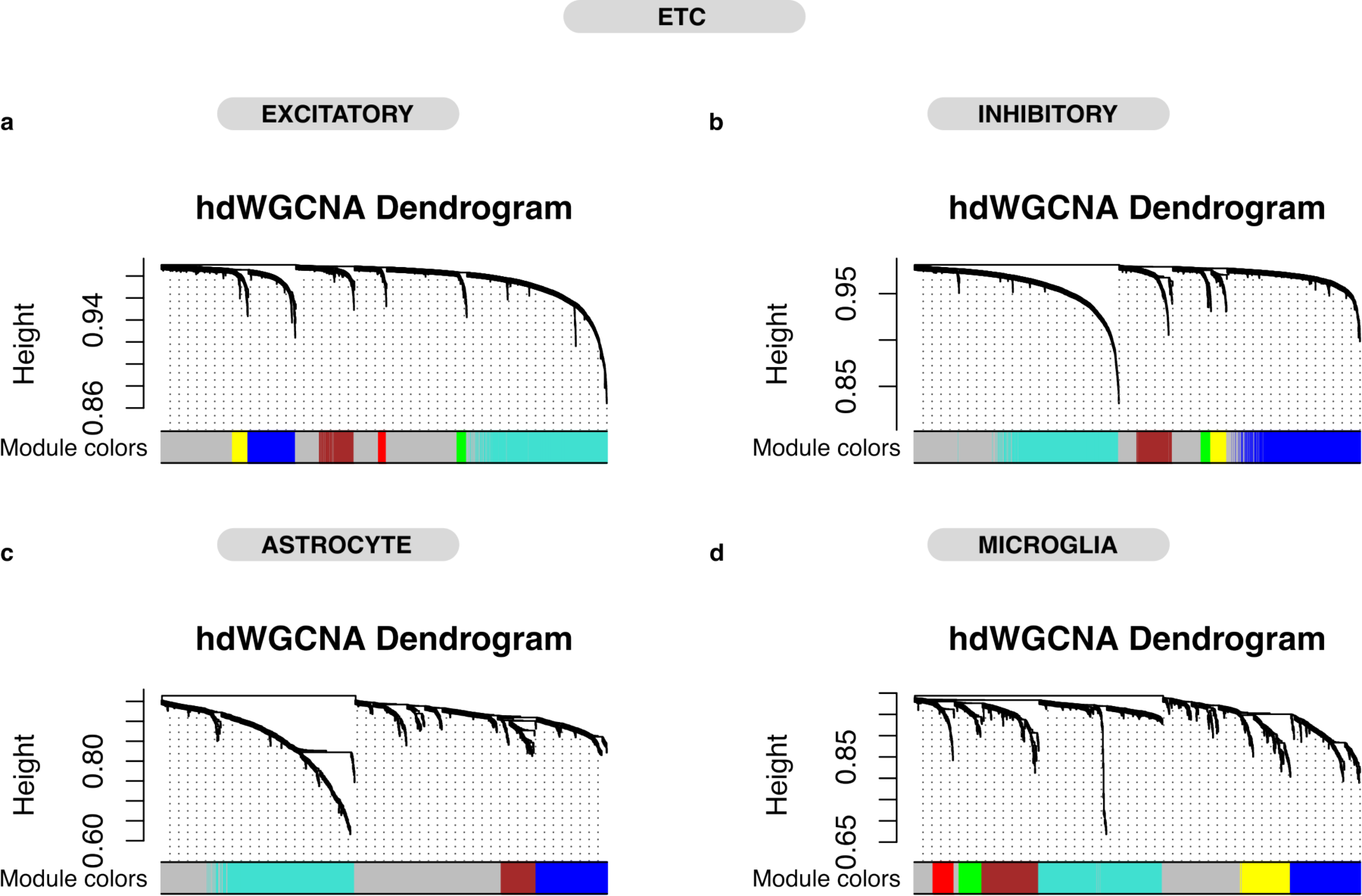

## References

1. Ferri CP, Prince M, Brayne C, Brodaty H, Fratiglioni L, Ganguli M, et al. Global prevalence of dementia: a Delphi consensus study. The Lancet. 2005 Dec 17;366(9503):2112–7.

2. Reitz C, Mayeux R. Alzheimer disease: Epidemiology, diagnostic criteria, risk factors and biomarkers. Biochemical Pharmacology. 2014 Apr 15;88(4):640–51.

3. Gouras GK, Tampellini D, Takahashi RH, Capetillo-Zarate E. Intraneuronal β-amyloid accumulation and synapse pathology in Alzheimer’s disease. Acta Neuropathol. 2010 May;119(5):523–41.

4. Hyman BT, Van Hoesen GW, Damasio AR, Barnes CL. Alzheimer’s Disease: Cell-Specific Pathology Isolates the Hippocampal Formation. Science. 1984 Sep 14;225(4667):1168–70.

5. Mondragón-Rodríguez S, Basurto-Islas G, Santa-Maria I, Mena R, Binder LI, Avila J, et al. Cleavage and conformational changes of tau protein follow phosphorylation during Alzheimer’s disease. International Journal of Experimental Pathology. 2008;89(2):81– 90.

6. Vogt LJK, Hyman BT, Van Hoesen GW, Damasio AR. Pathological alterations in the amygdala in Alzheimer’s disease. Neuroscience. 1990 Jan 1;37(2):377–85.

7. Gómez-Isla T, Hollister R, West H, Mui S, Growdon JH, Petersen RC, et al. Neuronal loss correlates with but exceeds neurofibrillary tangles in Alzheimer’s disease. Annals of Neurology. 1997;41(1):17–24.

8. Mucke L, Selkoe DJ. Neurotoxicity of Amyloid β-Protein: Synaptic and Network Dysfunction. Cold Spring Harb Perspect Med. 2012 Jul;2(7):a006338.

9. Parodi J, Sepúlveda FJ, Roa J, Opazo C, Inestrosa NC, Aguayo LG. β-Amyloid Causes Depletion of Synaptic Vesicles Leading to Neurotransmission Failure*. Journal of Biological Chemistry. 2010 Jan 22;285(4):2506–14.

10. Selkoe DJ. Alzheimer’s Disease Is a Synaptic Failure. Science. 2002 Oct 25;298(5594):789–91.

11. Wang M, Li A, Sekiya M, Beckmann ND, Quan X, Schrode N, et al. Molecular Networks and Key Regulators of the Dysregulated Neuronal System in Alzheimer’s Disease [Internet]. bioRxiv; 2019 [cited 2022 Sep 7]. p. 788323. Available from: https://www.biorxiv.org/content/10.1101/788323v1

12. Mrdjen D, Fox EJ, Bukhari SA, Montine KS, Bendall SC, Montine TJ. The basis of cellular and regional vulnerability in Alzheimer’s disease. Acta Neuropathol. 2019 Nov;138(5):729–49.

13. Wang X, Michaelis ML, Michaelis EK. Functional Genomics of Brain Aging and Alzheimer’s Disease: Focus on Selective Neuronal Vulnerability. Curr Genomics. 2010 Dec;11(8):618–33.

14. Crist AM, Hinkle KM, Wang X, Moloney CM, Matchett BJ, Labuzan SA, et al. Transcriptomic analysis to identify genes associated with selective hippocampal vulnerability in Alzheimer’s disease. Nature Communications. 2021;12(1).

15. Roussarie JP, Yao V, Rodriguez-Rodriguez P, Oughtred R, Rust J, Plautz Z, et al. Selective neuronal vulnerability in Alzheimer’s disease: a network-based analysis. Neuron. 2020 Sep 9;107(5):821–835.e12.

16. Stranahan AM, Mattson MP. Selective vulnerability of neurons in layer II of the entorhinal cortex during aging and Alzheimer’s disease. Neural Plast. 2010;2010:108190.

17. Wang M, Roussos P, McKenzie A, Zhou X, Kajiwara Y, Brennand KJ, et al. Integrative network analysis of nineteen brain regions identifies molecular signatures and networks underlying selective regional vulnerability to Alzheimer’s disease. Genome Medicine. 2016 Nov 1;8(1):104.

18. Cuevas-Diaz Duran R, González-Orozco JC, Velasco I, Wu JQ. Single-cell and single- nuclei RNA sequencing as powerful tools to decipher cellular heterogeneity and dysregulation in neurodegenerative diseases. Frontiers in Cell and Developmental Biology [Internet]. 2022 [cited 2023 Feb 8];10. Available from: https://www.frontiersin.org/articles/10.3389/fcell.2022.884748

19. Luquez T, Gaur P, Kosater IM, Lam M, Lee DI, Mares J, et al. Cell type-specific changes identified by single-cell transcriptomics in Alzheimer’s disease. Genome Medicine. 2022 Nov 30;14(1):136.

20. Leng K, Li E, Eser R, Piergies A, Sit R, Tan M, et al. Molecular characterization of selectively vulnerable neurons in Alzheimer’s disease. Nat Neurosci. 2021 Feb;24(2):276–87.

21. Mathys H, Davila-Velderrain J, Peng Z, Gao F, Mohammadi S, Young JZ, et al. Single- cell transcriptomic analysis of Alzheimer’s disease. Nature. 2019 Jun;570(7761):332–7.

22. Morabito S, Miyoshi E, Michael N, Shahin S, Martini AC, Head E, et al. Single-nucleus chromatin accessibility and transcriptomic characterization of Alzheimer’s disease. Nat Genet. 2021 Aug;53(8):1143–55.

23. Zhou Y, Song WM, Andhey PS, Swain A, Levy T, Miller KR, et al. Human and mouse single-nucleus transcriptomics reveal TREM2-dependent and TREM2-independent cellular responses in Alzheimer’s disease. Nat Med. 2020 Jan;26(1):131–42.

24. Grubman A, Chew G, Ouyang JF, Sun G, Choo XY, McLean C, et al. A single-cell atlas of entorhinal cortex from individuals with Alzheimer’s disease reveals cell-type-specific gene expression regulation. Nat Neurosci. 2019 Dec;22(12):2087–97.

25. Keren-Shaul H, Spinrad A, Weiner A, Matcovitch-Natan O, Dvir-Szternfeld R, Ulland TK, et al. A Unique Microglia Type Associated with Restricting Development of Alzheimer’s Disease. Cell. 2017 Jun 15;169(7):1276–1290.e17.

26. Habib N, McCabe C, Medina S, Varshavsky M, Kitsberg D, Dvir-Szternfeld R, et al. Disease-associated astrocytes in Alzheimer’s disease and aging. Nat Neurosci. 2020 Jun;23(6):701–6.

27. Castrillo JI, Oliver SG. Alzheimer’s as a Systems-Level Disease Involving the Interplay of Multiple Cellular Networks. Methods Mol Biol. 2016;1303:3–48.

28. Rayaprolu S, Higginbotham L, Bagchi P, Watson CM, Zhang T, Levey AI, et al. Systems-based proteomics to resolve the biology of Alzheimer’s disease beyond amyloid and tau. Neuropsychopharmacol. 2021 Jan;46(1):98–115.

29. Calabrò M, Rinaldi C, Santoro G, Crisafulli C. The biological pathways of Alzheimer disease: a review. AIMS Neurosci. 2020 Dec 16;8(1):86–132.

30. Guo T, Zhang D, Zeng Y, Huang TY, Xu H, Zhao Y. Molecular and cellular mechanisms underlying the pathogenesis of Alzheimer’s disease. Molecular Neurodegeneration. 2020 Jul 16;15(1):40.

31. Morabito S, Reese F, Rahimzadeh N, Miyoshi E, Swarup V. hdWGCNA identifies co- expression networks in high-dimensional transcriptomics data. Cell Reports Methods [Internet]. 2023 Jun 12 [cited 2023 Jun 13];0(0). Available from: https://www.cell.com/cell-reports-methods/abstract/S2667-2375(23)00127-3

32. Miyoshi E, Morabito S, Henningfield CM, Rahimzadeh N, Shabestari SK, Das S, et al. Spatial and single-nucleus transcriptomic analysis of genetic and sporadic forms of Alzheimer’s Disease [Internet]. bioRxiv; 2023 [cited 2023 Jul 27]. p. 2023.07.24.550282. Available from: https://www.biorxiv.org/content/10.1101/2023.07.24.550282v1

33. Langfelder P, Mischel PS, Horvath S. When Is Hub Gene Selection Better than Standard Meta-Analysis? PLOS ONE. 2013 Apr 17;8(4):e61505.

34. de la Fuente A. From ‘differential expression’ to ‘differential networking’ – identification of dysfunctional regulatory networks in diseases. Trends in Genetics. 2010 Jul 1;26(7):326–33.

35. Gabitto MI, Travaglini KJ, Rachleff VM, Kaplan ES, Long B, Ariza J, et al. Integrated multimodal cell atlas of Alzheimer’s disease [Internet]. bioRxiv; 2023 [cited 2023 May 9]. p. 2023.05.08.539485. Available from: https://www.biorxiv.org/content/10.1101/2023.05.08.539485v1

36. Hyman BT, Phelps CH, Beach TG, Bigio EH, Cairns NJ, Carrillo MC, et al. National Institute on Aging–Alzheimer’s Association guidelines for the neuropathologic assessment of Alzheimer’s disease. Alzheimers Dement. 2012 Jan;8(1):1–13.

37. Braak H, Braak E. Development of Alzheimer-related neurofibrillary changes in the neocortex inversely recapitulates cortical myelogenesis. Acta Neuropathol. 1996 Aug;92(2):197–201.

38. Thal DR, Rüb U, Orantes M, Braak H. Phases of Aβ-deposition in the human brain and its relevance for the development of AD. Neurology. 2002 Jun 25;58(12):1791–800.

39. Braak H, Alafuzoff I, Arzberger T, Kretzschmar H, Del Tredici K. Staging of Alzheimer disease-associated neurofibrillary pathology using paraffin sections and immunocytochemistry. Acta Neuropathol. 2006 Oct;112(4):389–404.

40. Grinberg LT, Ferretti RE de L, Farfel JM, Leite R, Pasqualucci CA, Rosemberg S, et al. Brain bank of the Brazilian aging brain study group - a milestone reached and more than 1,600 collected brains. Cell Tissue Bank. 2007;8(2):151–62.

41. Hänzelmann S, Castelo R, Guinney J. GSVA: gene set variation analysis for microarray and RNA-Seq data. BMC Bioinformatics. 2013 Jan 16;14(1):7.

42. SB. Synapse | Sage Bionetworks [Internet]. [cited 2024 Feb 22]. Available from: https://www.synapse.org

43. Seattle Alzheimer’s Disease Brain Cell Atlas (SEA-AD) - Registry of Open Data on AWS [Internet]. [cited 2024 Feb 22]. Available from: https://registry.opendata.aws/allen-sea-ad-atlas/

44. L. Lun AT, Bach K, Marioni JC. Pooling across cells to normalize single-cell RNA sequencing data with many zero counts. Genome Biology. 2016 Apr 27;17(1):75.

45. Seattle Alzheimer’s Disease Brain Cell Atlas - brain-map.org [Internet]. [cited 2024 Feb 22]. Available from: https://portal.brain-map.org/explore/seattle-alzheimers-disease/seattle-alzheimers-disease-brain-cell-atlas-download?edit&language=en

46. Squair JW, Gautier M, Kathe C, Anderson MA, James ND, Hutson TH, et al. Confronting false discoveries in single-cell differential expression. Nat Commun. 2021 Sep 28;12(1):5692.

47. Love MI, Huber W, Anders S. Moderated estimation of fold change and dispersion for RNA-seq data with DESeq2. Genome Biology. 2014 Dec 5;15(12):550.

48. Xie Z, Bailey A, Kuleshov MV, Clarke DJB, Evangelista JE, Jenkins SL, et al. Gene Set Knowledge Discovery with Enrichr. Current Protocols. 2021 Mar;1(3):e90.

49. Blanchard JW, Akay LA, Davila-Velderrain J, von Maydell D, Mathys H, Davidson SM, et al. APOE4 impairs myelination via cholesterol dysregulation in oligodendrocytes. Nature. 2022 Nov;611(7937):769–79.

50. Subramanian A, Tamayo P, Mootha VK, Mukherjee S, Ebert BL, Gillette MA, et al. Gene set enrichment analysis: A knowledge-based approach for interpreting genome- wide expression profiles. Proceedings of the National Academy of Sciences. 2005 Oct 25;102(43):15545–50.

51. Mohammadi S, Davila-Velderrain J, Kellis M. A multiresolution framework to characterize single-cell state landscapes. Nat Commun. 2020 Oct 26;11(1):5399.

52. Butler A, Hoffman P, Smibert P, Papalexi E, Satija R. Integrating single-cell transcriptomic data across different conditions, technologies, and species. Nat Biotechnol. 2018 May;36(5):411–20.

53. Ochoa D, Hercules A, Carmona M, Suveges D, Baker J, Malangone C, et al. The next- generation Open Targets Platform: reimagined, redesigned, rebuilt. Nucleic Acids Research. 2023 Jan 6;51(D1):D1353–9.

54. Kanehisa M, Furumichi M, Sato Y, Kawashima M, Ishiguro-Watanabe M. KEGG for taxonomy-based analysis of pathways and genomes. Nucleic Acids Res. 2023 Jan 6;51(D1):D587–92.

55. Rouillard AD, Gundersen GW, Fernandez NF, Wang Z, Monteiro CD, McDermott MG, et al. The harmonizome: a collection of processed datasets gathered to serve and mine knowledge about genes and proteins. Database. 2016 Jan 1;2016:baw100.

56. Law CW, Zeglinski K, Dong X, Alhamdoosh M, Smyth GK, Ritchie ME. A guide to creating design matrices for gene expression experiments [Internet]. F1000Research; 2020 [cited 2023 Mar 17]. Available from: https://f1000research.com/articles/9-1444

57. Papalexi E, Satija R. Single-cell RNA sequencing to explore immune cell heterogeneity. Nat Rev Immunol. 2018 Jan;18(1):35–45.

58. Skinnider MA, Squair JW, Foster LJ. Evaluating measures of association for single-cell transcriptomics. Nat Methods. 2019 May;16(5):381–6.

59. Chen S, Mar JC. Evaluating methods of inferring gene regulatory networks highlights their lack of performance for single cell gene expression data. BMC Bioinformatics. 2018 Jun 19;19(1):232.

60. van Dam S, Võsa U, van der Graaf A, Franke L, de Magalhães JP. Gene co-expression analysis for functional classification and gene–disease predictions. Briefings in Bioinformatics. 2018 Jul 20;19(4):575–92.

61. Langfelder P, Horvath S. WGCNA: an R package for weighted correlation network analysis. BMC Bioinformatics. 2008 Dec 29;9(1):559.

62. Manczak M, Park BS, Jung Y, Reddy PH. Differential expression of oxidative phosphorylation genes in patients with Alzheimer’s disease: implications for early mitochondrial dysfunction and oxidative damage. Neuromolecular Med. 2004;5(2):147– 62.

63. Berridge MJ. Calcium hypothesis of Alzheimer’s disease. Pflugers Arch. 2010 Feb;459(3):441–9.

64. Liu L, Wu Q, Zhong W, Chen Y, Zhang W, Ren H, et al. Microarray Analysis of Differential Gene Expression in Alzheimer’s Disease Identifies Potential Biomarkers with Diagnostic Value. Med Sci Monit. 2020 Jan 27;26:e919249-1-e919249-16.

65. Morabito S, Miyoshi E, Michael N, Swarup V. Integrative genomics approach identifies conserved transcriptomic networks in Alzheimer’s disease. Human Molecular Genetics. 2020 Oct 10;29(17):2899–919.

66. Workgroup AACH, Khachaturian ZS. Calcium Hypothesis of Alzheimer’s disease and brain aging: A framework for integrating new evidence into a comprehensive theory of pathogenesis. Alzheimer’s & Dementia. 2017;13(2):178–182.e17.

67. Ceglia I, Reitz C, Gresack J, Ahn JH, Bustos V, Bleck M, et al. APP intracellular domain/WAVE1 pathway reduces amyloid β production. Nat Med. 2015 Sep;21(9):1054–9.

68. Ochaba J, Monteys AM, O’Rourke JG, Reidling JC, Steffan JS, Davidson BL, et al. PIAS1 regulates mutant Huntingtin accumulation and Huntington’s disease-associated phenotypes in vivo. Neuron. 2016 May 4;90(3):507–20.

69. He K, Zhang J, Liu J, Cui Y, Liu LG, Ye S, et al. Functional genomics study of protein inhibitor of activated STAT1 in mouse hippocampal neuronal cells revealed by RNA sequencing. Aging (Albany NY). 2021 Mar 24;13(6):9011–27.

70. Anderson AG, Rogers BB, Loupe JM, Rodriguez-Nunez I, Roberts SC, White LM, et al. Single nucleus multiomics identifies ZEB1 and MAFB as candidate regulators of Alzheimer’s disease-specific cis-regulatory elements. Cell Genomics. 2023 Feb 2;100263.

71. Bohush A, Bieganowski P, Filipek A. Hsp90 and Its Co-Chaperones in Neurodegenerative Diseases. Int J Mol Sci. 2019 Oct 9;20(20):4976.

72. Gonzalez-Rodriguez M, Villar-Conde S, Astillero-Lopez V, Villanueva-Anguita P, Ubeda-Banon I, Flores-Cuadrado A, et al. Neurodegeneration and Astrogliosis in the Human CA1 Hippocampal Subfield Are Related to hsp90ab1 and bag3 in Alzheimer’s Disease. Int J Mol Sci. 2021 Dec 23;23(1):165.

73. Labbadia J, Morimoto RI. The biology of proteostasis in aging and disease. Annu Rev Biochem. 2015;84:435–64.

74. Miron J, Picard C, Labonté A, Auld D, Breitner J, Poirier J, et al. Association of PPP2R1A with Alzheimer’s disease and specific cognitive domains. Neurobiol Aging. 2019 Sep;81:234–43.

75. Del Prete D, Checler F, Chami M. Ryanodine receptors: physiological function and deregulation in Alzheimer disease. Mol Neurodegener. 2014 Jun 1;9:21.

76. Yao J, Sun B, Institoris A, Zhan X, Guo W, Song Z, et al. Limiting RyR2 Open Time Prevents Alzheimer’s Disease-Related Neuronal Hyperactivity and Memory Loss but Not β-Amyloid Accumulation. Cell Rep. 2020 Sep 1;32(12):108169.

77. Yao J, Liu Y, Sun B, Zhan X, Estillore JP, Turner RW, et al. Increased RyR2 open probability induces neuronal hyperactivity and memory loss with or without Alzheimer’s disease-causing gene mutations. Alzheimers Dement. 2022 Nov;18(11):2088–98.

78. Ashraf A, Jeandriens J, Parkes HG, So PW. Iron dyshomeostasis, lipid peroxidation and perturbed expression of cystine/glutamate antiporter in Alzheimer’s disease: Evidence of ferroptosis. Redox Biol. 2020 May 1;32:101494.

79. Foster EM, Dangla-Valls A, Lovestone S, Ribe EM, Buckley NJ. Clusterin in Alzheimer’s Disease: Mechanisms, Genetics, and Lessons From Other Pathologies. Frontiers in Neuroscience [Internet]. 2019 [cited 2023 Aug 14];13. Available from: https://www.frontiersin.org/articles/10.3389/fnins.2019.00164

80. Lazarev VF, Tsolaki M, Mikhaylova ER, Benken KA, Shevtsov MA, Nikotina AD, et al. Extracellular GAPDH Promotes Alzheimer Disease Progression by Enhancing Amyloid-β Aggregation and Cytotoxicity. Aging Dis. 2021 Aug 1;12(5):1223–37.

81. Tang L, Wang ZB, Ma LZ, Cao XP, Tan L, Tan MS. Dynamic changes of CSF clusterin levels across the Alzheimer’s disease continuum. BMC Neurol. 2022 Dec 1;22(1):508.

82. Chen T, Gai WP, Abbott CA. Dipeptidyl peptidase 10 (DPP10(789)): a voltage gated potassium channel associated protein is abnormally expressed in Alzheimer’s and other neurodegenerative diseases. Biomed Res Int. 2014;2014:209398.

83. Malamon JS, Kriete A. Erosion of Gene Co-expression Networks Reveal Deregulation of Immune System Processes in Late-Onset Alzheimer’s Disease. Frontiers in Neuroscience [Internet]. 2020 [cited 2023 Jul 21];14. Available from: https://www.frontiersin.org/articles/10.3389/fnins.2020.00228

84. Mitra S, P KB, R SC, Saikumar NV, Philip P, Narayanan M. Alzheimer’s disease rewires gene coexpression networks coupling different brain regions [Internet]. bioRxiv; 2022 [cited 2023 Jul 7]. p. 2022.05.22.492888. Available from: https://www.biorxiv.org/content/10.1101/2022.05.22.492888v1

85. Xiang J, Wang X, Gao Y, Li T, Cao R, Yan T, et al. Phosphodiesterase 4D Gene Modifies the Functional Network of Patients With Mild Cognitive Impairment and Alzheimer’s Disease. Front Genet. 2020 Jan 1;11:890.

86. Tibbo AJ, Tejeda GS, Baillie GS. Understanding PDE4’s function in Alzheimer’s disease; a target for novel therapeutic approaches. Biochem Soc Trans. 2019 Oct 31;47(5):1557–65.

87. Shi Y, Lv J, Chen L, Luo G, Tao M, Pan J, et al. Phosphodiesterase-4D Knockdown in the Prefrontal Cortex Alleviates Memory Deficits and Synaptic Failure in Mouse Model of Alzheimer’s Disease. Front Aging Neurosci. 2021 Jan 1;13:722580.

88. Qiang Q, Skudder-Hill L, Toyota T, Wei W, Adachi H. CSF GAP-43 as a biomarker of synaptic dysfunction is associated with tau pathology in Alzheimer’s disease. Sci Rep. 2022 Oct 1;12(1):17392.

89. Sandelius Å, Portelius E, Källén Å, Zetterberg H, Rot U, Olsson B, et al. Elevated CSF GAP-43 is Alzheimer’s disease specific and associated with tau and amyloid pathology. Alzheimers Dement. 2019 Jan 1;15(1):55–64.

90. Zhu Y, Guo X, Zhu F, Zhang Q, Yang Y, For TADNI. Association of CSF GAP-43 and *APOE* ε4 with Cognition in Mild Cognitive Impairment and Alzheimer’s Disease. Cells. 2022 Dec 1;12(1):13.

91. Fernandez-Enright F, Andrews JL. Lingo-1: a novel target in therapy for Alzheimer’s disease? Neural Regen Res. 2016 Jan;11(1):88–9.

92. Xiang Y, Xin J, Le W, Yang Y. Neurogranin: A Potential Biomarker of Neurological and Mental Diseases. Frontiers in Aging Neuroscience [Internet]. 2020 [cited 2023 Aug 16];12. Available from: https://www.frontiersin.org/articles/10.3389/fnagi.2020.584743

93. Folts CJ, Giera S, Li T, Piao X. Adhesion G protein-coupled receptors as drug target for neurological diseases. Trends Pharmacol Sci. 2019 Apr;40(4):278–93.

94. Kulczyńska-Przybik A, Dulewicz M, Słowik A, Borawska R, Kułakowska A, Kochanowicz J, et al. The Clinical Significance of Cerebrospinal Fluid Reticulon 4 (RTN4) Levels in the Differential Diagnosis of Neurodegenerative Diseases. J Clin Med. 2021 Nov 13;10(22):5281.

95. Zhang H, Therriault J, Kang MS, Ng KP, Pascoal TA, Rosa-Neto P, et al. Cerebrospinal fluid synaptosomal-associated protein 25 is a key player in synaptic degeneration in mild cognitive impairment and Alzheimer’s disease. Alzheimer’s Research & Therapy. 2018 Aug 16;10(1):80.

96. Zolochevska O, Bjorklund N, Woltjer R, Wiktorowicz JE, Taglialatela G. Postsynaptic Proteome of Non-Demented Individuals with Alzheimer’s Disease Neuropathology. J Alzheimers Dis. 2018 Jan 1;65(2):659–82.

97. Han J, Hyun J, Park J, Jung S, Oh Y, Kim Y, et al. Aberrant role of pyruvate kinase M2 in the regulation of gamma-secretase and memory deficits in Alzheimer’s disease. Cell Rep. 2021 Dec 7;37(10):110102.

98. Li Y, Chen Z, Wang Q, Lv X, Cheng Z, Wu Y, et al. Identification of hub proteins in cerebrospinal fluid as potential biomarkers of Alzheimer’s disease by integrated bioinformatics. J Neurol. 2023 Mar 1;270(3):1487–500.

99. Traxler L, Herdy JR, Stefanoni D, Eichhorner S, Pelucchi S, Szücs A, et al. Warburg- like metabolic transformation underlies neuronal degeneration in sporadic Alzheimer’s disease. Cell Metab. 2022 Sep 1;34(9):1248–1263.e6.

100. Zhou X, Sun L, Bastos de Oliveira F, Qi X, Brown WJ, Smolka MB, et al. Prosaposin facilitates sortilin-independent lysosomal trafficking of progranulin. J Cell Biol. 2015 Sep 1;210(6):991–1002.

101. Mendsaikhan A, Tooyama I, Bellier JP, Serrano GE, Sue LI, Lue LF, et al. Characterization of lysosomal proteins Progranulin and Prosaposin and their interactions in Alzheimer’s disease and aged brains: increased levels correlate with neuropathology. Acta Neuropathologica Communications. 2019 Dec 21;7(1):215.

102. Zhang Q, Ma C, Gearing M, Wang PG, Chin LS, Li L. Integrated proteomics and network analysis identifies protein hubs and network alterations in Alzheimer’s disease. Acta Neuropathologica Communications. 2018 Mar 1;6(1):19.

103. Anirudhan A, Angulo-Bejarano PI, Paramasivam P, Manokaran K, Kamath SM, Murugesan R, et al. RPL6: A Key Molecule Regulating Zinc- and Magnesium-Bound Metalloproteins of Parkinson’s Disease. Front Neurosci. 2021 Mar 11;15:631892.

104. Pollutri D, Penzo M. Ribosomal Protein L10: From Function to Dysfunction. Cells. 2020 Nov 19;9(11):2503.

105. Garcia-Esparcia P, Diaz-Lucena D, Ainciburu M, Torrejón-Escribano B, Carmona M, Llorens F, et al. Glutamate Transporter GLT1 Expression in Alzheimer Disease and Dementia With Lewy Bodies. Front Aging Neurosci. 2018 Jan 1;10:122.

106. Yeung JHY, Palpagama TH, Wood OWG, Turner C, Waldvogel HJ, Faull RLM, et al. EAAT2 Expression in the Hippocampus, Subiculum, Entorhinal Cortex and Superior Temporal Gyrus in Alzheimer’s Disease. Front Cell Neurosci. 2021 Jan 1;15:702824.

107. Chaudhury AR, Gerecke KM, Wyss JM, Morgan DG, Gordon MN, Carroll SL. Neuregulin-1 and erbB4 immunoreactivity is associated with neuritic plaques in Alzheimer disease brain and in a transgenic model of Alzheimer disease. J Neuropathol Exp Neurol. 2003 Jan 1;62(1):42–54.

108. León A, Aparicio GI, Scorticati C. Neuronal Glycoprotein M6a: An Emerging Molecule in Chemical Synapse Formation and Dysfunction. Front Synaptic Neurosci. 2021 May 4;13:661681.

109. Liu F, Gong X, Yao X, Cui L, Yin Z, Li C, et al. Variation in the CACNB2 gene is associated with functional connectivity of the Hippocampus in bipolar disorder. BMC Psychiatry. 2019 Feb 11;19(1):62.

110. Woo RS, Lee JH, Yu HN, Song DY, Baik TK. Expression of ErbB4 in the apoptotic neurons of Alzheimer’s disease brain. Anat Cell Biol. 2010 Dec 1;43(4):332–9.

111. Woo RS, Lee JH, Yu HN, Song DY, Baik TK. Expression of ErbB4 in the neurons of Alzheimer’s disease brain and APP/PS1 mice, a model of Alzheimer’s disease. Anat Cell Biol. 2011 Jun 1;44(2):116–27.

112. Hüttenrauch M, Ogorek I, Klafki H, Otto M, Stadelmann C, Weggen S, et al. Glycoprotein NMB: a novel Alzheimer’s disease associated marker expressed in a subset of activated microglia. Acta Neuropathol Commun. 2018 Oct 1;6(1):108.

113. Satoh J ichi, Kino Y, Yanaizu M, Ishida T, Saito Y. Microglia express GPNMB in the brains of Alzheimer’s disease and Nasu-Hakola disease. Intractable Rare Dis Res. 2019 May;8(2):120–8.

114. Smith AM, Davey K, Tsartsalis S, Khozoie C, Fancy N, Tang SS, et al. Diverse human astrocyte and microglial transcriptional responses to Alzheimer’s pathology. Acta Neuropathol. 2022 Jan 1;143(1):75–91.

115. Zhu Z, Liu Y, Li X, Zhang L, Liu H, Cui Y, et al. GPNMB mitigates Alzheimer’s disease and enhances autophagy via suppressing the mTOR signal. Neurosci Lett. 2022 Jan 1;767:136300.

116. Ando K, Nagaraj S, Küçükali F, de Fisenne MA, Kosa AC, Doeraene E, et al. PICALM and Alzheimer’s Disease: An Update and Perspectives. Cells. 2022 Dec 10;11(24):3994.

117. Xu W, Tan L, Yu JT. The Role of PICALM in Alzheimer’s Disease. Mol Neurobiol. 2015 Aug 1;52(1):399–413.

118. Cimino PJ, Sokal I, Leverenz J, Fukui Y, Montine TJ. DOCK2 is a microglial specific regulator of central nervous system innate immunity found in normal and Alzheimer’s disease brain. Am J Pathol. 2009 Oct;175(4):1622–30.

119. Cimino PJ, Yang Y, Li X, Hemingway JF, Cherne MK, Khademi SB, et al. Ablation of the Microglial Protein DOCK2 Reduces Amyloid Burden in a Mouse Model of Alzheimer’s Disease. Exp Mol Pathol. 2013 Apr;94(2):366–71.

120. Öhrfelt A, Brinkmalm A, Dumurgier J, Brinkmalm G, Hansson O, Zetterberg H, et al. The pre-synaptic vesicle protein synaptotagmin is a novel biomarker for Alzheimer’s disease. Alzheimers Res Ther. 2016 Oct 3;8(1):41.

121. Mann CN, Shreedarshanee SD, Kersting CT, Bleem AV, Karch CM, Holtzman DM, et al. Astrocytic α2-Na+/K+ ATPase inhibition suppresses astrocyte reactivity and reduces neurodegeneration in a tauopathy mouse model. Sci Transl Med. 2022 Feb 16;14(632):eabm4107.

122. Robinson SR. Neuronal expression of glutamine synthetase in Alzheimer’s disease indicates a profound impairment of metabolic interactions with astrocytes. Neurochem Int. 2000 Apr 1;36(4–5):471–82.

123. Mendsaikhan A, Tooyama I, Serrano GE, Beach TG, Walker DG. Loss of Lysosomal Proteins Progranulin and Prosaposin Associated with Increased Neurofibrillary Tangle Development in Alzheimer Disease. J Neuropathol Exp Neurol. 2021 Aug 10;80(8):741–53.

124. Haydon PG, Carmignoto G. Astrocyte Control of Synaptic Transmission and Neurovascular Coupling. Physiological Reviews. 2006 Jul;86(3):1009–31.

125. Lian H, Zheng H. Signaling pathways regulating neuron–glia interaction and their implications in Alzheimer’s disease. Journal of Neurochemistry. 2016;136(3):475–91.

126. Stobart J, Anderson C. Multifunctional role of astrocytes as gatekeepers of neuronal energy supply. Frontiers in Cellular Neuroscience [Internet]. 2013 [cited 2023 Aug 31];7. Available from: https://www.frontiersin.org/articles/10.3389/fncel.2013.00038

127. Iannuzzi F, Sirabella R, Canu N, Maier TJ, Annunziato L, Matrone C. Fyn Tyrosine Kinase Elicits Amyloid Precursor Protein Tyr682 Phosphorylation in Neurons from Alzheimer’s Disease Patients. Cells. 2020 Jul 1;9(8):E1807.

128. Lau DHW, Hogseth M, Phillips EC, O’Neill MJ, Pooler AM, Noble W, et al. Critical residues involved in tau binding to fyn: implications for tau phosphorylation in Alzheimer’s disease. Acta Neuropathol Commun. 2016 May 1;4(1):49.

129. Gourmaud S, Paquet C, Dumurgier J, Pace C, Bouras C, Gray F, et al. Increased levels of cerebrospinal fluid JNK3 associated with amyloid pathology: links to cognitive decline. J Psychiatry Neurosci. 2015 May;40(3):151–61.

130. Musi CA, Agrò G, Santarella F, Iervasi E, Borsello T. JNK3 as Therapeutic Target and Biomarker in Neurodegenerative and Neurodevelopmental Brain Diseases. Cells. 2020 Sep 28;9(10):2190.

131. Solas M, Vela S, Smerdou C, Martisova E, Martínez-Valbuena I, Luquin MR, et al. JNK Activation in Alzheimer’s Disease Is Driven by Amyloid β and Is Associated with Tau Pathology. ACS Chem Neurosci. 2023 Apr 19;14(8):1524–34.

132. De Schepper S, Ge JZ, Crowley G, Ferreira LSS, Garceau D, Toomey CE, et al. Perivascular cells induce microglial phagocytic states and synaptic engulfment via SPP1 in mouse models of Alzheimer’s disease. Nat Neurosci. 2023 Mar;26(3):406–15.

133. Seaman MNJ, Mukadam AS, Breusegem SY. Inhibition of TBC1D5 activates Rab7a and can enhance the function of the retromer cargo-selective complex. J Cell Sci. 2018 Jun 1;131(12):jcs217398.

134. Barthelson K, Newman M, Lardelli M. Sorting Out the Role of the *Sortilin-Related Receptor 1* in Alzheimer’s Disease. J Alzheimers Dis Rep. 2020 May 1;4(1):123–40.

135. Chen SM, Yi YL, Zeng D, Tang YY, Kang X, Zhang P, et al. Hydrogen Sulfide Attenuates β2-Microglobulin-Induced Cognitive Dysfunction: Involving Recovery of Hippocampal Autophagic Flux. Front Behav Neurosci. 2019 Jan 1;13:244.

136. Ciarlo E, Massone S, Penna I, Nizzari M, Gigoni A, Dieci G, et al. An intronic ncRNA- dependent regulation of SORL1 expression affecting Aβ formation is upregulated in post-mortem Alzheimer’s disease brain samples. Dis Model Mech. 2013 Mar 1;6(2):424–33.

137. Gaiser AK, Bauer S, Ruez S, Holzmann K, Fändrich M, Syrovets T, et al. Serum Amyloid A1 Induces Classically Activated Macrophages: A Role for Enhanced Fibril Formation. Front Immunol. 2021 Jan 1;12:691155.

138. Hung C, Tuck E, Stubbs V, van der Lee SJ, Aalfs C, van Spaendonk R, et al. SORL1 deficiency in human excitatory neurons causes APP-dependent defects in the endolysosome-autophagy network. Cell Rep. 2021 Jun 1;35(11):109259.

139. Malik BR, Maddison DC, Smith GA, Peters OM. Autophagic and endo-lysosomal dysfunction in neurodegenerative disease. Molecular Brain. 2019 Nov 29;12(1):100.

140. Dai DL, Li M, Lee EB. Human Alzheimer’s disease reactive astrocytes exhibit a loss of homeostastic gene expression. Acta Neuropathologica Communications. 2023 Aug 2;11(1):127.

141. Jin J, Liu L, Chen W, Gao Q, Li H, Wang Y, et al. The Implicated Roles of Cell Adhesion Molecule 1 (CADM1) Gene and Altered Prefrontal Neuronal Activity in Attention-Deficit/Hyperactivity Disorder: A “Gene–Brain–Behavior Relationship”? Front Genet. 2019 Sep 26;10:882.

142. Gomez AM, Traunmüller L, Scheiffele P. Neurexins: molecular codes for shaping neuronal synapses. Nat Rev Neurosci. 2021 Mar;22(3):137–51.

143. Stogsdill JA, Ramirez J, Liu D, Kim YH, Baldwin KT, Enustun E, et al. Astrocytic neuroligins control astrocyte morphogenesis and synaptogenesis. Nature. 2017 Nov 8;551(7679):192–7.

144. Chung WS, Welsh CA, Barres BA, Stevens B. Do Glia Drive Synaptic and Cognitive Impairment in Disease? Nat Neurosci. 2015 Nov;18(11):1539–45.

145. Yu Y, Chen R, Mao K, Deng M, Li Z. The Role of Glial Cells in Synaptic Dysfunction: Insights into Alzheimer’s Disease Mechanisms. Aging Dis. 2023 Jul 26;

146. Henstridge CM, Tzioras M, Paolicelli RC. Glial Contribution to Excitatory and Inhibitory Synapse Loss in Neurodegeneration. Frontiers in Cellular Neuroscience [Internet]. 2019 [cited 2023 Sep 4];13. Available from: https://www.frontiersin.org/articles/10.3389/fncel.2019.00063

147. Braak H, Braak E. Neuropathological stageing of Alzheimer-related changes. Acta Neuropathol. 1991 Sep 1;82(4):239–59.

148. Braak H, Braak E. Staging of alzheimer’s disease-related neurofibrillary changes. Neurobiology of Aging. 1995 May 1;16(3):271–8.

